# Biomolecular models of EPI-X4 binding to CXCR4 allow the rational optimization of peptides with therapeutic potential

**DOI:** 10.1101/2020.10.23.352708

**Authors:** Pandian Sokkar, Mirja Harms, Christina Stürzel, Andrea Gilg, Gönül Kizilsavas, Martina Raasholm, Nico Preising, Manfred Wagner, Ludger Ständker, Gilbert Weidinger, Jan Münch, Elsa Sanchez-Garcia

**Affiliations:** Computational Biochemistry, Center of Medical Biotechnology, University of Duisburg-Essen, Essen, Germany; Faculty of Allied Health Science, Chettinad Hospital and Research Institute, Chettinad Academy of Research and Education, Kelambakkam, Tamil Nadu, India; Institute of Molecular Virology, Ulm University Medical Center, Ulm, Germany; Max Planck Institute for Polymer Research, Mainz, Germany; Institute of Biochemistry and Molecular Biology, Ulm University, Ulm, 89081, Germany; Core Facility Functional Peptidomics, Ulm University Medical Center, Ulm, 89081, Germany

**Keywords:** CXCR4, EPI-X4, molecular dynamics, rational design, peptide inhibitors

## Abstract

The Endogenous Peptide Inhibitor of CXCR4 (EPI-X4) is a body-own fragment of albumin and specific antagonist of the CXC-motif-chemokine receptor 4 (CXCR4). CXCR4 signaling is induced by its sole chemokine ligand CXCL12 and is involved in a plethora of functions including cell homing, differentiation, survival and angiogenesis. Consequently, dysregulation of CXCR4 is involved in a variety of disorders, such as cancer or inflammatory diseases, making CXCR4 an attractive drug target. EPI-X4 and derivatives with increased CXCR4 binding affinities represent promising leads as CXCR4 antagonists and have shown therapeutic activity in mouse models of inflammatory diseases. However, it is currently unclear how EPI-X4 and its derivatives interact with CXCR4. Here, by combining biomolecular simulations with experimental mutagenesis and activity studies we investigated the binding behavior of EPI-X4 to CXCR4 at the molecular level. Our work allowed us to show that the EPI-X4 peptide interacts primarily in the minor pocket of CXCR4 through its N-terminal residues. The biomolecular interactions highlighted by the computational studies are in good agreement with the experimental mutagenesis data. Moreover, we found that the N-terminal seven amino-acids of EPI-X4 (a 16-mer) and its improved derivatives (12-mers) are sufficient for CXCR4 binding, which led to the development of shorter leads with optimized CXCR4 antagonizing properties. Collectively, we here established how EPI-X4 binds to its receptor and used this knowledge for rational drug design. The new peptide variants developed by us are more potent in terms of inhibiting CXCR4-downstream signaling and cancer cell migration, without toxic effects.

## INTRODUCTION

Chemokine receptors are important mediators of numerous processes in the human body. Among them is C-X-C motif chemokine receptor 4 (CXCR4), a 365-residue rhodopsin-like G-protein coupled receptor (GPCR). CXCR4 is expressed on ubiquitous hematopoietic and non-hematopoietic tissues where it regulates important processes, such as immune response, development, hematopoiesis, vascularization, tissue renewal and regeneration.^1^ It is therefore not surprising that the faulty regulation of CXCR4 is responsible for several pathologies, i.e. inflammatory diseases,^2^ immunodeficiencies or cancer.^3,4^ In many different forms of cancer, CXCR4 is often overexpressed or overactivated,^1,5^ which is linked to cancer progression by promoting proliferation, survival and metastasis.^6,7^ CXCR4 expressing cells migrate in direction of CXCL12, the sole endogenous chemokine ligand of CXCR4. CXCL12 is mainly expressed in the bone marrow, the lymph nodes, lung, and liver; tissues where primary metastasis mainly occur.^8^ Besides that, CXCR4 is a major entry coreceptor of HIV-1.^9^ Thus, CXCR4 represents a promising drug target. In the last years, intense research was performed to identify and develop CXCR4 antagonists for therapeutic applications in HIV/AIDS, cancer and inflammatory disorders. However, the only FDA-approved CXCR4 antagonist so far is AMD3100, which is restricted to single treatments due to its severe side effects.^10^

Because of the importance of CXCR4 in physiological and pathological processes, crystal structures of CXCR4 in complex with different ligands have been determined; (i) with the viral chemokine antagonist vMIP-II,^11^ (ii) with the small molecule antagonist IT1t,^12^ and (iii) with the cyclic CVX15 peptide (analogue of polyphemusin).^12^ As a member of the GPCR family, the structure of CXCR4 consists of a canonical bundle of seven transmembrane (TM) α-helices, three intracellular (ICL) and three extracellular (ECL) loops. The extracellular N-terminus of CXCR4 features a 34-residue intrinsically disordered loop that forms a disulfide bridge with C274 of helix VII through C28. CXCR4 has a relatively large and open binding pocket that is located at the extracellular region.^8,12^ This binding pocket is defined by the seven TM domains and is decorated by negatively charged aspartate and glutamate residues.^11^ It can be separated into loosely defined major and minor subpockets; the first comprised of TMs III, IV, V, VI and VII and the later comprised of TMs I, II, III and VII.^13^ While in GPCRs most ligands only interact with the major subpocket, for CXCR4 it was shown that the small molecule antagonist IT1t,^12^ as well as the chemokines vMIP-II and CXCL12,^11^ interact with the minor subpocket. In case of CXCL12, the N-terminus of CXCR4 mediates binding and is responsible for receptor activation.^11,14^ K1_CXCL12_ interacts with D97_CXCR4_ and E288_CXCR4_ through the N-terminal amine group of CXCL12. Another residue that is thought to mediate receptor activation is D187_CXCR4_ (ECL2) that interacts with S4_CXCL12_ and Y7_CXCL12_.^14^

Zirafi et. al. identified the endogenous CXCR4 antagonist EPI-X4 (Endogenous Peptide Inhibitor of CXCR4) by screening a peptide library derived from human hemofiltrate.^15,16^ They found that this 16-mer peptide is derived by proteolytic degradation of human serum albumin by pH-regulated proteases, e.g. cathepsin D, and is therefore endogenously present at acidic sites of the body. EPI-X4 binds specifically to CXCR4 and interrupts the interaction of CXCL12 with its receptor thereby antagonizing chemokine-mediated effects, like cell migration and infiltration. Additionally, EPI-X4 has inverse antagonistic properties, since it reduces basal receptor signaling activity. So far, the physiological function of EPI-X4 is not clear, however, the peptide was found in the urine of patients with renal failure and therefore might have regulatory functions in the body.^16^

Optimized analogues of EPI-X4 include the C-terminally truncated 12-mer derivative WSC02 that harbors 4 amino acid substitutions compared to the wild type (L1I, Y4W, T5S, Q10C). WSC02 has about 30-fold increased potency compared to the wild type peptide.^16^ A further optimized 12-mer version, EPI-X4 JM#21, harbors three additional amino acid substitutions (V2L, K6R, V9L), which led to further increase in receptor affinity (Harms et al. preprint).^17^ In addition, EPI-X4 JM#21 showed increased efficiency for inhibition of CXCL12-mediated receptor signaling (> 100-fold compared to EPI-X4, 10-fold compared to WSC02) and cancer cell migration (~ 1,500-fold compared to EPI-X4, 30-fold compared to WSC02). Also, optimized EPI-X4 derivatives showed very promising therapeutic effects in mouse models for stem cell mobilization, allergic airway eosinophilia and atopic dermatitis.^16,17^

Computational modeling has also been carried out to determine the interaction sites of different CXCR4 receptor ligands with the binding pockets of CXCR4.^18,19^ Several molecular dynamics (MD) studies are reported, primarily (i) to characterize the structure and function of CXCR4 and agonist binding,^20–22^ (ii) to study the interactions between CXCR4 and small molecule/peptide antagonists,^23–25^ and (iii) to study the dimerization of CXCR4^26–28^. However, the interactions of EPI-X4 with CXCR4 have so far not been investigated in detail.

Here, we used extensive biomolecular simulations, mutagenesis and activity studies to identify the interaction sites of the endogenous peptide EPI-X4 with CXCR4, in order to better understand its function and effects on the receptor. The study of the interaction motifs of the optimized EPI-X4 derivatives WSC02 and JM#21 with CXCR4 also provides insights for peptide optimization. Thus, the knowledge provided by the analysis of the biomolecular simulations of these peptides with the receptor allowed us to design shorter EPI-X4 variants in which the activity of the parent compound is preserved, while improving their potential for therapeutic applications. The design of shorter active peptides is a key step towards biomedical applications. These reduced variants have the advantage of synthetic accessibility and lower production costs, reduced dosage, and open the opportunity for oral delivery.

## RESULTS AND DISCUSSION

### Binding modes of EPI-X4 to CXCR4

We used various computational approaches to determine the binding mode of EPI-X4 to CXCR4, explicitly considering solvation and the membrane environment (Figure 1a). First, we performed docking calculations with the complete structure of CXCR4 (PDB IDs: 3ODU and 2K04, see computational details)^12,29^ using three different conformations of EPI-X4 taken from the previously published NMR ensemble (2N0X)^16^. We also modeled one binding mode based on the reported structure of the complex of CXCR4 with the vMIP-II peptide (4RWS)^11^. For the ease of discussion, these four binding poses are abbreviated as MID-IN1, MID-IN2, CTER-IN and NTER-IN (Figure 1b-e). Although there is a large amount of conformations possible for this system, these models provided a comprehensive starting point for molecular dynamics (MD) simulations assessing the interactions between CXCR4 and EPI-X4.

**Figure 1.**
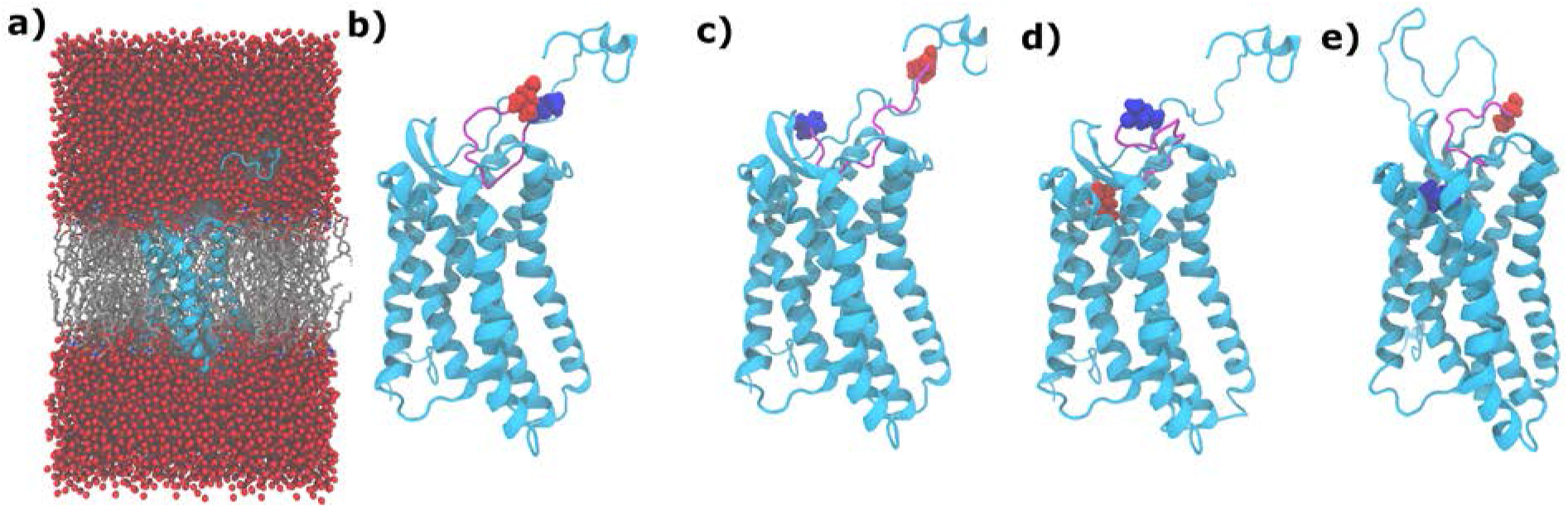
Binding modes and MD simulations setup of the CXCR4/EPI-X4 complex. CXCR4 (cyan) and EPI-X4 (purple) are shown in cartoon representation and the N-terminal (blue) and C-terminal (red) residues of EPI-X4 are shown as spheres **a**) Representative all-atom CXCR4/EPI-X4 complex embedded in a POPC (1-palmitoyl-2-oleoyl-sn-glycero-3-phosphocholine) lipid bilayer (grey sticks) and water (red spheres) as used in the MD simulations. Bilayer and water are omitted in the other figures of this manuscript for clarity. The binding modes represented in **b**, **c** and **d** were obtained using docking calculations and the binding mode shown in **e** was obtained using homology modeling. In **b**) and **c**) the middle portion of the peptide is inside the binding pocket (abbreviated as MID-IN1 and MID-IN2, respectively. In **d**) the C-terminus of the peptide is inside the binding pocket (CTER-IN) while in **e**), the N-terminus of EPI-X4 is the region inside the binding pocket (NTER-IN).

### Coarse-Grained (CG) simulations

Independently of the all-atom MD simulations that will be described on the next section, we investigated the self-assembly of the CXCR4/EPI-X4 complex from the unbound state. To this end, we used steered MD simulations at the coarse-grained resolution (MARTINI 2.2 CG force field)^30^. The coarse-grained system comprised the receptor, peptide, water, ions, and lipid bilayer. For the sake of simplicity, we used a derivative of EPI-X4 comprising the nine N-terminal residues (408-416). We applied biasing forces to the peptide on three different points (i.e., residues 1, 5 and 9) towards the binding pocket. The pulling forces were equal in magnitude (same force constant) on all the three sites and were operating simultaneously. This ensures that the pulling is not biased to a particular site on the peptide. The aim of these simulations was to determine if there is a preferred pattern or binding mode. Therefore, we used very small force constants to allow plenty of conformation sampling that reduces the bias associated with initial configurations. Using twenty different initial structures, we performed twenty simulations, each for 1 μs and under three force constants setups: 0.1, 0.2 and 0.5 kJ/mol/Å^2^. Our results indicated that when the force constant is very small (*k_f_*=0.1 kJ/mol/Å^2^ and *k_f_*=0.2 kJ/mol/Å^2^), 6/20 trajectories resulted in the formation of NTER-IN complexes (Figure 2). For the rest of the trajectories, the peptide could not enter the binding pocket. In other words, there is a certain amount of energetic barrier to overcome the steric and/or electrostatic repulsions involved in the binding process. When the force constant was large enough (*k_f_*=0.5 kJ/mol/Å^2^), 14/20 trajectories resulted in the NTER-IN binding mode and 1/20 in CTER-IN. Thus, these pulling simulations indicate that NTER-IN is favored over the other modes.

**Figure 2.**
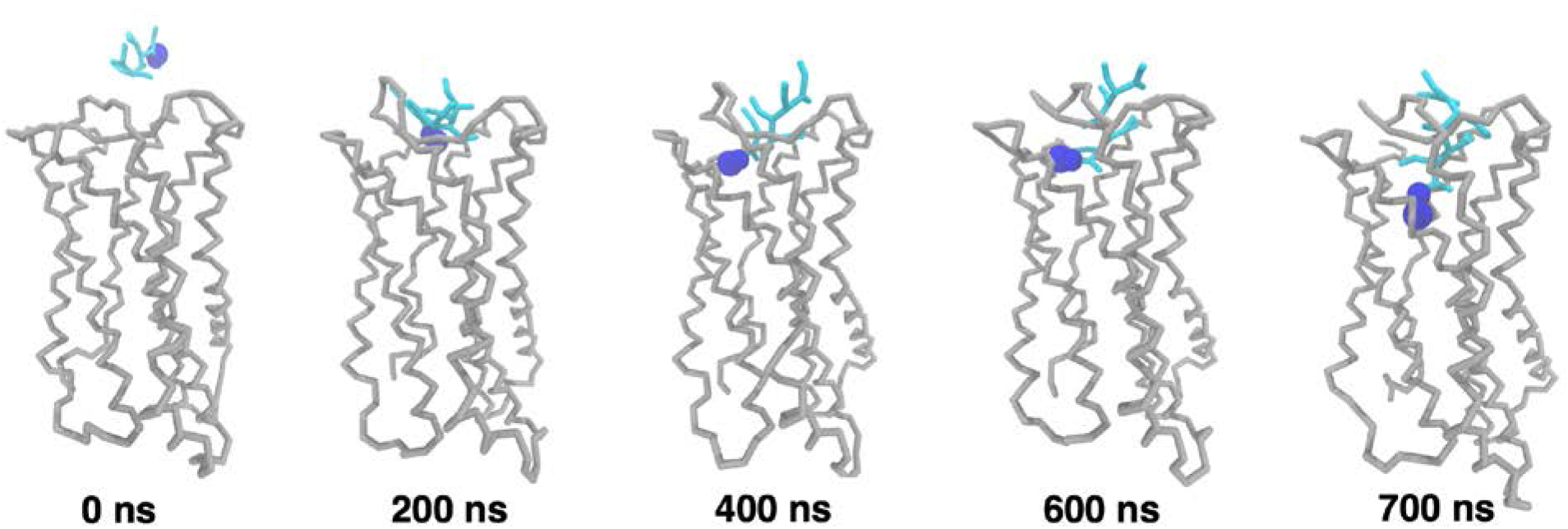
Snapshots of a CG MD simulation with pulling force (*k_f_*=0.2 kJ/mol/Å^2^). CXCR4 is shown as grey sticks and the peptide is shown as cyan sticks. The L1 residue is indicated with blue spheres to highlight the N-terminal region of EPI-X4.

### All-atom molecular dynamics simulations

Owing to the inherent approximations of CG models, these simulations do not provide detailed information at the molecular level of the binding interactions between CXCR4 and EPI-X4. Therefore, we performed all-atom MD simulations of the four systems (MID-IN1, MID-IN2, CTER-IN and NTER-IN, Figure 1), three replicas in each case. We used full atom resolution, including explicit solvent and the lipid bilayer (Figure 1a). The analysis of the MDs indicated that the CTER-IN and NTER-IN motifs showed the lowest amount of structural variations in most of the trajectories, also suggesting that these two binding modes are favorable (Figure S1, Supplementary computational details). Therefore, we extended the simulation time for CTER-IN and NTER-IN from 600 ns to 1.35 μs each. The NTER-IN binding mode showed negligible fluctuation in residues 1-8 (Figure S1c), which are known to be critical for the binding of EPI-X4 derivatives.^16^

The solvent accessible surface area (SASA) of the peptide during the MD simulations (Table 1, Figure S2) indicates more solvent exposure of the peptide in the cases of MID-IN1 and MID-IN2, whereas in the CTER-IN and NTER-IN modes, EPI-X4 is buried slightly deeper into the receptor (Table 1). In addition, the larger protein-peptide interaction interface in the CTER-IN and NTER-IN motifs suggests that the interaction of EPI-X4 with CXCR4 is stronger in these cases with respect to MID-IN1 and MID-IN2.

**Table 1.** SASA and protein-peptide interaction interface area of EPI-X4 in the different binding modes^a^

| Binding mode | SASA (Å<sup>2</sup>) | Protein-peptide<br>Interface area (Å<sup>2</sup>) |
| --- | --- | --- |
| MID-IN1 | 1329 | 810 |
| MID-IN2 | 1470 | 836 |
| CTER-IN | 1053<br>(1072) | 1061<br>(991) |
| NTER-IN | 1159<br>(1098) | 1103<br>(1137) |

<sup>a</sup> We also show in parenthesis data from 600 ns simulations of CTER-IN and NTER-IN for comparison using the same simulation time in all four cases.

Next, we carried out a clustering analysis of the simulation trajectories (Figure 3). The MID-IN2 motif did not produce any meaningfully populated cluster (5% for the highest populated cluster). MID-IN1 resulted in a cluster of conformations 22% populated in ~600 ns simulation (Figure 3a). CTER-IN exhibited two different clusters, both of which are significantly populated (32% and 31%) in 1.35 μs. The difference between these two clusters is that in cluster#1 (Figure 3b) EPI-X4 interacts in the “major binding pocket” of CXCR4, whereas in cluster#2 (Figure 3c) EPI-X4 partially occupies both the “minor” and “major” binding pockets of the receptor. The most interesting observation was in the case of the NTER-IN mode, which shows the highest population of the conformations (70%) in a single cluster in 1.35 μs. As seen in Figure 3d, the N-terminal region of EPI-X4 interacts with the “minor” pocket of CXCR4, in a similar manner to the vMIP-II peptide.^11^ We note that the same tendencies were observed if only 600 ns trajectories of CTER-IN and NTER-IN were analyzed (i.e., in CTER-IN 31% of each cluster and in NTER-IN 86% of the structures were in a single cluster).

**Figure 3.**
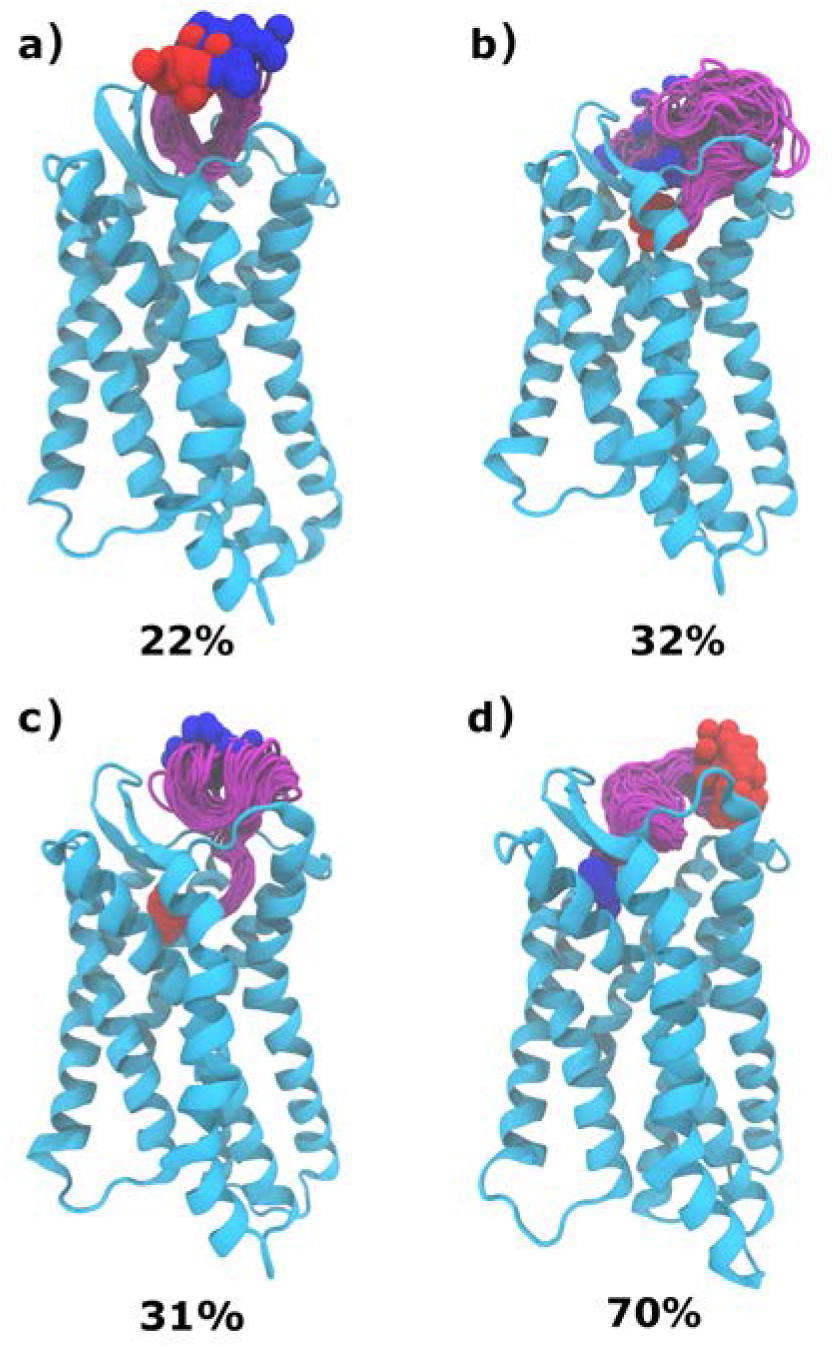
Clustering analysis from the MD simulations of a) MID-IN1, b and c) CTER-IN, and d) NTER-IN. The color scheme is the same as in Figure 1. Structures of the same cluster are shown to illustrate the spread of conformations. The clustering analysis was performed with a RMSD cutoff of 3 Å using the VMD program. ^31^

Next, we focused on the specific interactions that stabilize the complexes of EPI-X4 and CXCR4. We calculated the number of hydrogen bonds (averaged over the simulation time) formed between CXCR4 and EPI-X4 during the MD simulations. The NTER-IN binding motif exhibited the highest number of hydrogen bonding (H-bond) interactions with an average of 5 H-bonds. MID -IN 1 and MID-IN2 featured a smaller amount of H-bond contacts (average of 2.6 and 3.1 H-bonds, respectively). By comparison, CTER-IN shows a slightly higher number of H-bonds (average 3.8). This result agrees with the previous findings (RMSD, SASA, peptide-protein surface and clustering analysis) that indicate that MID-IN and MID-IN2 are not favored binding modes when compared to the CTER-IN and NTER-IN complexes.

As mentioned, NTER-IN shows the highest population of H-bonding contacts (Table S1). The H-bond involving T5_EPI-X4_ and D187_CXCR4_ sidechains was found to be populated by 69.5% during the 1.35 μs simulation of the NTER-IN mode. Similarly, H-bonds involving L1 (mainchain), R3, T5, K6 and K7 were also found to be significantly populated in this mode. This suggests that these residues are playing a pivotal role in the interaction with the predominantly negatively charged binding pocket of CXCR4.

In addition, the analysis of the total interaction energies and the corresponding van der Waals (vdW) and electrostatic contributions indicates that NTER-IN exhibits high interaction strength compared to the other binding modes, followed by CTER-IN (Table S2). Interestingly, the vdW contribution to the interaction energy is the same in NTER-IN and CTER-IN. Thus, the difference in their interaction strength is mainly due to electrostatic contributions. This trend is in good agreement with the previously discussed results, all of them pointing to NTER-IN as the most favorable binding mode.

Next, we decomposed the interaction energies by residue-wise contributions (Figure 4). Our analysis indicates that positively charged residues provide the largest favorable contribution to the interaction energies. This is because the binding pocket of CXCR4 is rich in negatively charged residues. Thus, the positively charged N-terminal Leu (L1) of EPI-X4 stabilizes the binding, whereas the negatively charged C-terminal Leu (L16) has a destabilizing effect. The contributions to the interaction energy of the positive charged residues L1, R3, K6 and K7 of EPI-X4 are nearly the same in all binding modes. Consequently, the improved binding affinity of the NTER-IN interaction motif may be related to the optimal positioning of the N-terminal residues V2, Y4 and T5.

**Figure 4.**
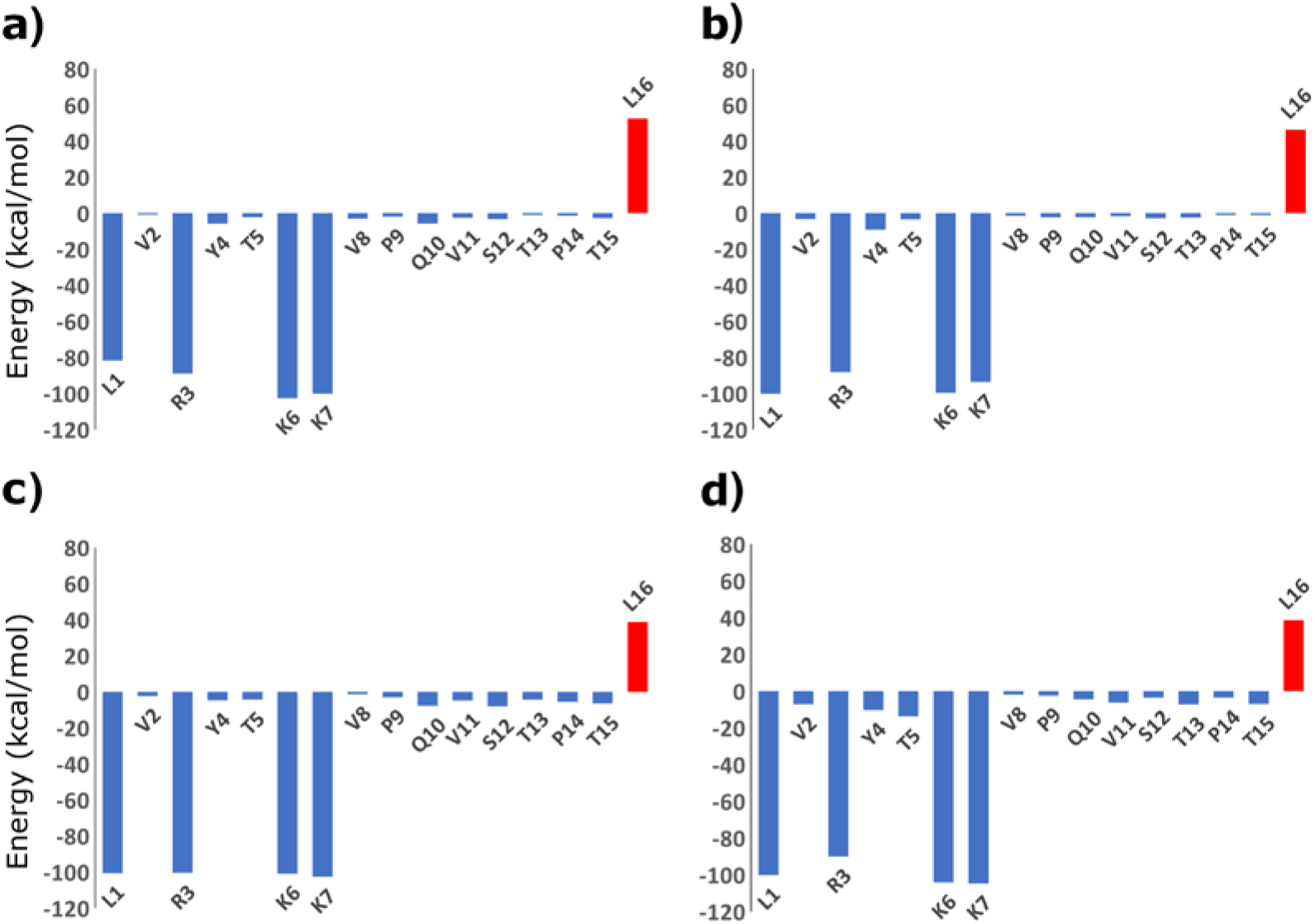
Contributions to the interaction energy by individual residues of EPI-X4 from the simulations of A) MID-IN, B) MID-IN2, C) CTER-IN and D) NTER-IN modes.

### EPI-X4/CXCR4 interaction sites

Previously, Zirafi et al. reported several EPI-X4 derivatives and their IC_50_ values. The authors showed that N-terminal residues of EPI-X4 are crucial for binding/inhibition of CXCR4.^16^ Mutations of Leu1 (L1A, L1G and L1F) or deletion of this amino acid render the peptide inactive. On the contrary, truncations at the C-terminus of up to 4 residues did not seem to influence CXCR4 binding or inhibition. It was also shown that the deletion of C-terminal residues Q10-L16 did not affect the binding drastically.

Our structural model of NTER-IN shows that the side chain of L1 fits effortlessly in the hydrophobic pocket area (Figure 5c and d), which allows rationalizing the lack of activity of the N-terminally mutations above-mentioned. With respect to EPI-X4, the L1F derivative has a larger and planar phenyl ring that would be sterically hindered in the hydrophobic pocket of CXCR4. On the contrary, L1A and L1G, that render the peptide inactive, have very small sidechains that cannot establish optimal contacts in such pocket.

**Figure 5.**
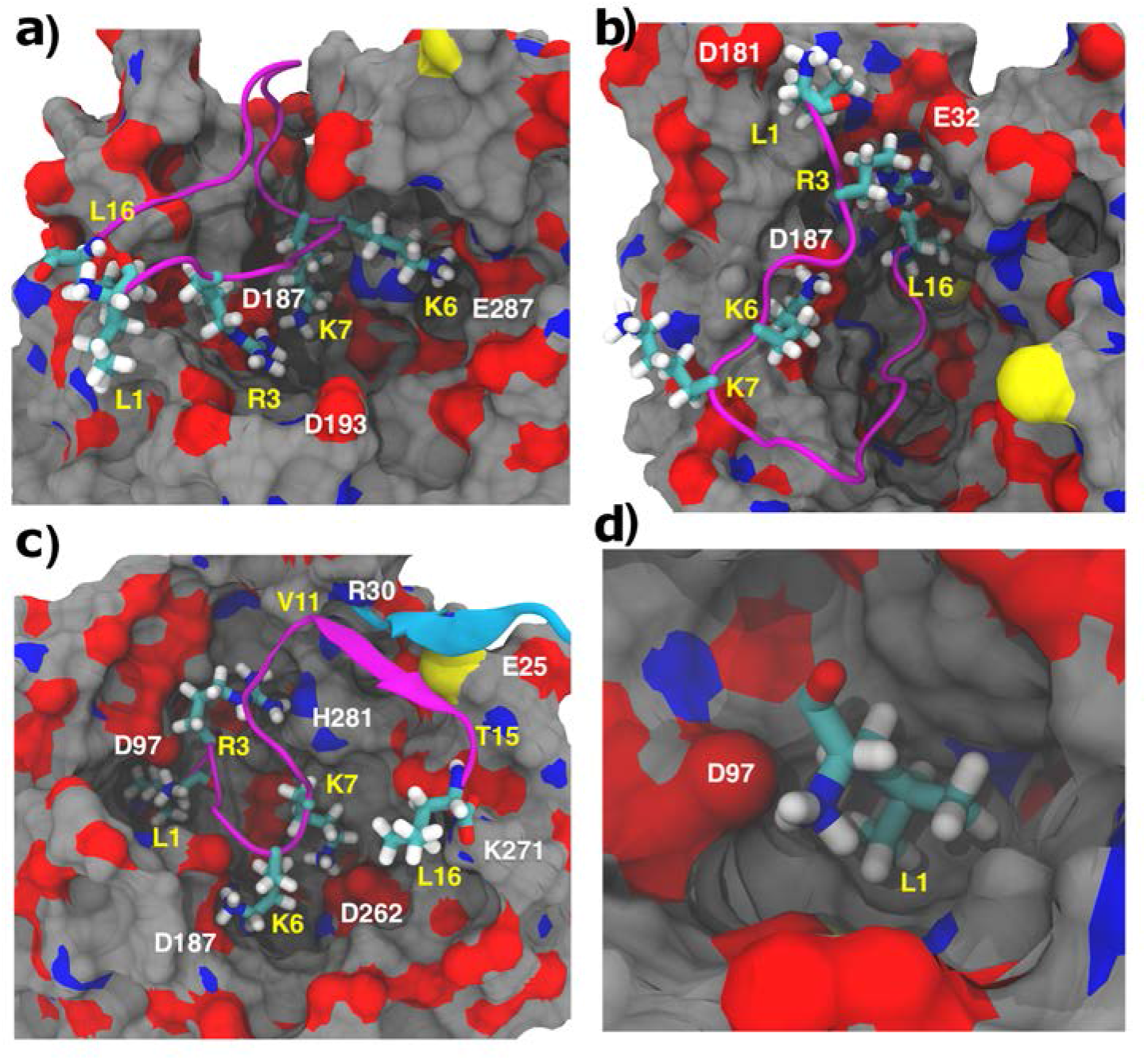
Binding modes MID-IN1 (a), CTER-IN (b) and NTER-IN (c, d) identified from the clustering analysis. CXCR4 is represented as surface (carbon = grey, oxygen = red and nitrogen = blue) and EPI-X4 is shown in cartoon diagram (purple). Important residues of EPI-X4 are shown as sticks and labelled in yellow. CXCR4 residues involved in the binding are labelled in white. The β-strand of CXCR4 comprising residues E25 – R30 is shown in cyan.

Unlike NTER-IN, the binding modes MID-IN, MID-IN2 and CTER-IN do not agree with the experimental findings stated above. This is because, in these three models, the N-terminal region of the peptide is exposed to the solvent (Figures 1 and 5). However, L1 was found to contribute to the stabilization of the complex in all binding modes (Figure 4). We decomposed the L1 interaction energy to identify the *sidechain* contribution, which was found to be 0.0, − 0.7 and −1.3 kcal/mol, for MID-IN, MID-IN2 and CTER-IN, respectively. For NTER-IN, the hydrophobic L1 sidechain contributes −6.5 kcal/mol to the interaction energy, indicating that the sidechain of L1 is optimally placed in terms of both electrostatic and vdW interactions.

In NTER-IN, the sidechain of L1_EPI-X4_ is buried in the hydrophobic cavity of the “minor” binding pocket of CXCR4 (formed by F93, W94, W102, V112 and Y116), while the backbone ammonium group of EPI-X4 is forming a salt-bridge with D97 of CXCR4 (Figure 5d). This arrangement also allows the peptide to establish a variety of interactions, such as the R3_EPI-X4_ – H281_CXCR4_ hydrogen bond, the salt bridges K6_EPI-X4_ – D187_CXCR4_, K7_EPI-X4_ – D262_CXCR4_ and L16(C-terminal)_EPI-X4_ – K271_CXCR4_ (Table S1). Another interesting aspect of this binding mode is the additional stabilization provided by the short β-strand involving residues V11 - Thr15 of EPI-X4, which interacts with the β-strand of CXCR4 comprising residues E25 - R30 (Figure 5c). These interactions were either absent or non-conserved in the other binding modes.

### Site-directed mutagenesis experiments

To further assess these findings, we performed site-directed mutagenesis experiments of CXCR4 residues involved in the interaction with EPI-X4. For this, we inserted single point mutations in the amino acid sequence of CXCR4 and transfected the receptor in 293T cells that have a very low endogenous expression level of wild type CXCR4. Transfected cells were then analyzed for CXCR4 surface expression (Figure S3) and EPI-X4 binding in an antibody competition assay.^32^ This assay is based on the competition of the monoclonal CXCR4 antibody 12G5 (an antibody that binds to a region close to the receptor orthosteric binding pocket) and CXCR4 ligands for receptor interaction.^32^ Mutations of residues predicted to be important for EPI-X4 binding strongly decreased receptor binding affinity (increase of 50% inhibitory concentration, IC_50_, Figures 6a and b, Table 2). When D97_CXCR4_ was replaced by uncharged residues (D97S, D97T, D97N, and D97Q), the binding was nearly abolished, indicating that this residue is essential for binding of EPI-X4. As predicted, the D97E mutation causes reduction of the binding affinity of EPI-X4 to CXCR4 (~20 fold), despite the high similarity between the amino acids Asp and Glu. This can be related to the slightly extended (by one –CH_2_– group) sidechain of Glu, which weakens the salt bridge with L1_EPI-X4_ and/or the hydrophobic interaction of the later with V112_CXCR4_, since these two interactions are coupled. Accordingly, V112A leads to only a small change in the IC_50_ values, whereas V112L causes a nearly 6-fold increase in the IC_50_ values, indicating that the smaller amino acids (Val or Ala) in this binding pocket are tolerated for the interaction with the sidechain of L1_EPI-X4_ unlike the larger L112 residue. Replacement of D187 by an amino acid with the same charge (D187E) has low or no effect in EPI-X4 binding. Interestingly, the D262A mutation did not lead to a loss of activity. There, we can speculate that the conformational flexibility associated to the introduction of the smaller alanine residue results in larger structural rearrangements allowing new stabilizing interactions between the peptide and the receptor. Furthermore, unlike D187 and D97, D262 seems to play a less prominent role in the interactions with the peptide (Table S1). Another mutation, L41A, results in a nearly 20-fold increase in IC_50_. In the NTER-IN model, the hydrophobic L41 sidechain is packed with the hydrophobic region of V2EPI-X4, suggesting that this interaction is relevant for the formation of the CXCR4/EPI-X4 complex. Therefore, it may be expected that a double mutation involving L41 (V112A+L41A) or a triple mutation (D97E+V112A+L41A) would increase the IC_50_ further. In contrast, these mutations seem to have recovered the loss of L41A activity. In case of the double mutant, the smaller sidechain of A112 offers flexibility to the interactions involving the L1_EPI-X4_ residue, which in turn allows V2_EPI-X4_ to establish contacts with the V41A sidechain. The flexibility of L1_EPI-X4_ might increase in the triple mutant, where the D97E sidechain is slightly extended, compared to the wild type. These synergistic effects could explain the trends observed in the L41A mutations. Detailed computational studies of the CXCR4 mutants are needed to fully rationalize the effect of such modifications on the interaction networks involved in ligand binding. Nevertheless, the mutagenesis experiments (Table 2) corroborated our computational results indicating that the first seven amino acids of EPI-X4 are essential for binding and that NTER-IN is the favored binding mode of EPI-X4 in CXCR4.

**Figure 6.**
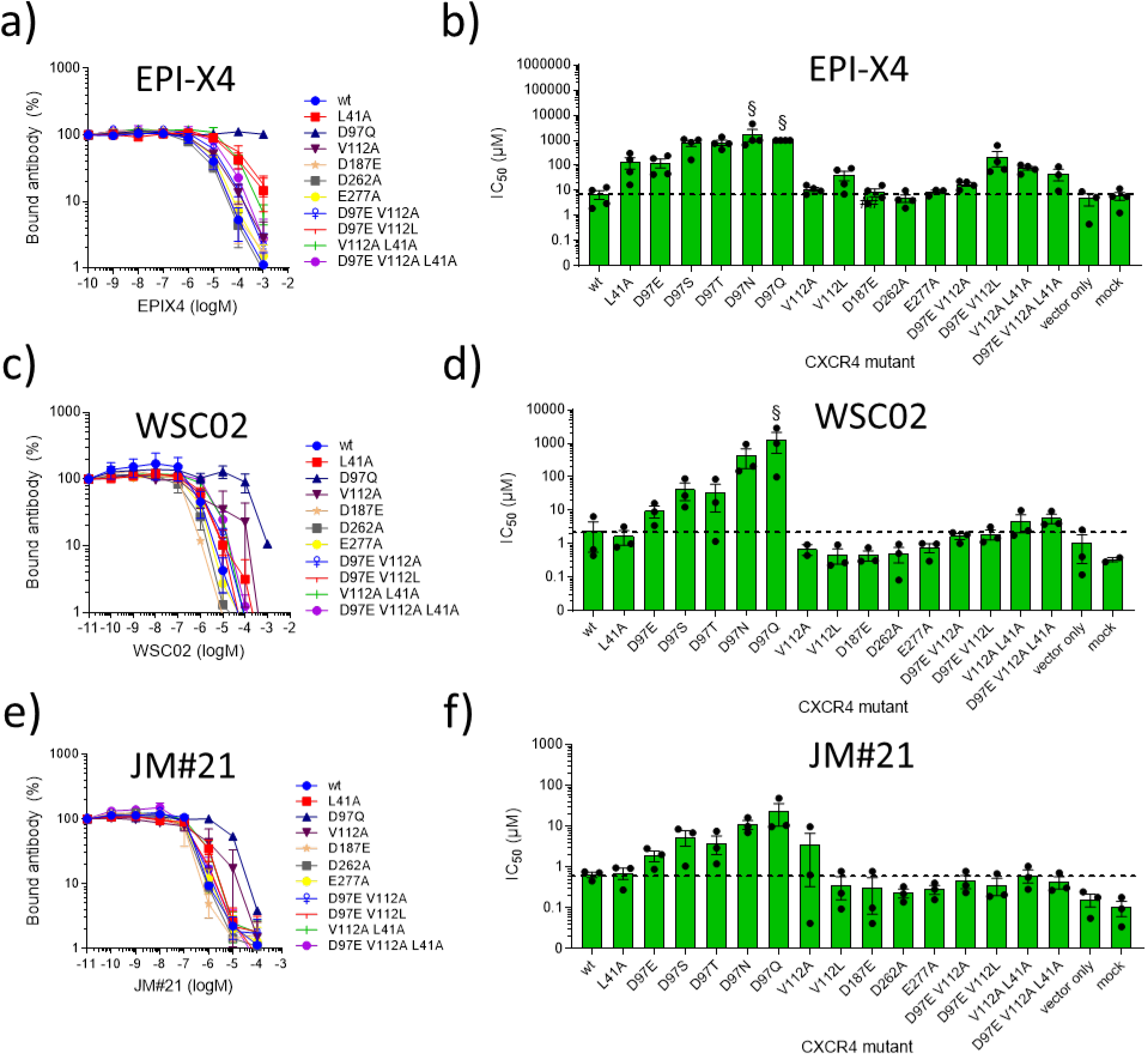
EPI-X4, and EPI-X4 derivatives WSC02 and JM#21 interactions with point-mutated CXCR4 as assessed by antibody competition. Amino acid substitutions were introduced in the sequence of CXCR4 by site-directed mutagenesis, cloned into an IRES-eGFP expression vector and transfected into 293T cells. Afterwards, cells were incubated with serially diluted EPI-X4 (a, b), WSC02 (c, d), or JM#21 (e, f) in presence of a constant concentration of CXCR4 specific antibody (clone 12G5). After 2 hours, bound antibody was analyzed by flow cytometry. a, c, and e) Representative dose dependent replacement of 12G5 antibody by peptides (See also Supplementary Figure S6). b, d and f) IC_50_ values were calculated by non-linear regression. Shown are data derived from at least 3 individual experiments ± SEM. § average IC_50_ of 1 or more values exceed 1000 μM.

**Table 2.** IC_50_ values of EPI-X4 determined in an 12G5-competition assay for CXCR4 mutants

| CXCR4 mutation | IC<sub>50</sub> ± SEM (μM) | Interaction with EPI-X4<sup>a</sup> | Agreement<sup>b</sup> |
| --- | --- | --- | --- |
| **wt** | **7.16 ± 2.78** | - |  |
| **L41A** | 136.98 ± 65.31 | V2 sidechain, hydrophobic | yes |
| **D97E** | 129.09 ± 49.75 | L1 N-terminal, salt bridge | yes |
| **D97S** | 780.18 ± 212.66 | L1 N-terminal, salt bridge | yes |
| **D97T** | 822.33 ± 197.45 | L1 N-terminal, salt bridge | yes |
| **D97N** | > 1000 | L1 N-terminal, salt bridge | yes |
| **D97Q** | > 1000 | L1 N-terminal, salt bridge | yes |
| **V112A** | 11.20 ± 2.00 | L1 sidechain, hydrophobic | yes |
| **V112L** | 41.62 ± 17.04 | L1 sidechain, hydrophobic | yes |
| **D187E** | 8.49 ± 2.87 | T5 sidechain, H-bonds | yes |
| **D262A** | 5.03 ± 1.69 | K7 sidechain, salt bridge | no |
| **E277A** | 8.77 ± 1.40 | T5 sidechain, H-bonds | yes |
| **D97E+V112A** | 17.90 ± 2.96 | - | yes |
| **D97E+V112L** | 221.40 ± 234.11 | - | yes |
| **V112A+L41A** | 77.41 ± 13.26 |  | yes |
| **D97E+V112A+L41A** | 46.92 ± 23.04 |  | yes |

<sup>a</sup>EPI-X4 residues interacting with wildtype residues of CXCR4 in the NTER-IN model
<sup>b</sup>Agreement between the NTER-IN model and the mutagenesis experiments

### Analysis of optimized EPI-X4 derivatives WSC02 and JM#21

Next, we investigated the interactions of the optimized EPI-X4 derivatives WSC02 and JM#21, that showed potent therapeutic effects in mouse models of atopic dermatitis and asthma ^16,17^. Both, WSC02 and JM#21, are C-terminally truncated analogues of EPI-X4 harboring 12 amino acids each. WSC02 is an optimized derivative with 4 amino acid substitutions (L1I, Y4W, T5S, Q10C) and has about 30-fold increased activity compared to its precursor. JM#21 is an optimized variant of WSC02 with three additional amino acid substitutions (V2L, K6R, V8L) and is about 35-fold more active than WSC02.^17^ First, we performed MD simulations of these peptides to compare their interaction patterns with respect to EPI-X4. Given the structural similarities between the peptides^17^, the binding modes of WSC02 and JM#21 were generated by homology modeling using the NTER-IN motif. In both cases, the complexes were subjected to 600 ns MD simulations (three replicas of 200 ns each).

Like in EPI-X4, the RMSD analysis indicated that the structure of CXCR4 is conserved during the simulations (Figures S4, a and c). Although for the peptides the RMSD values indicated structural fluctuations (Figure S4, b and d), these are largely focused in the C-terminal region (residues 7-12) of the peptides, which rearranges from the initial conformation, whereas the N-terminal segment (residues 1-6) did not change with respect to the initial pose. This conformational flexibility of the C-terminal region was also observed in the simulations of EPI-X4.

The clustering analysis indicated for both, WSC02/CXCR4 and JM#21/CXCR4, two predominant clusters of structures (populations of 29%,18% and 39%,16%, respectively), which differ only in the orientation of the C-terminal region of the peptide (Figure S5). We found that, with respect to the parental peptide and WSC02, JM#21 exhibited a somewhat larger amount of conserved hydrogen bonds. According to the model, R6_JM#21_ interacts with D262_CXCR4_ during 81% of the simulation time (Table S3). WSC02 and JM#21 both form H-bonds with E288_CXCR4_ through their N-terminal amino acid I1 (Table S3 and Figure 7), which was confirmed experimentally by mutagenesis analysis (Figure 6c-f and Table S3). Substitution of E288 to Ala or Asp nearly abolished binding of WSC02 and JM#21 to CXCR4. A hydrogen bond is also established between I1_WSC02_ and D97_CXCR4_ (Table S3). In our experimental setup, substitution of D97 strongly decreased binding of the peptides to CXCR4, similar to the results obtained with EPI-X4. Overall, WSC02 and JM#21 bind to CXCR4 like EPI-X4 through its N-terminus (Figure 7). However, WSC02 and JM#21 show similar interaction patterns involving their residues S5, K6/R6, and K7. In the three peptides, the interaction of T5 or S5 with D187_CXCR4_ is conserved. Experimentally, substitution of D187 by Glu did not have any significant influence on peptide binding. Substitution of D187 to Ala lead to interruption of receptor expression and, thus, could not be tested. The most notable difference among the peptides, WSC02 and JM#21, is the formation of a strong bidentate H-bond between R6_JM21_ and D262_CXCR4_. This could explain the improved binding affinity of JM#21 compared to WSC02 and EPI-X4. The analysis of the interaction energies indicates similar tendencies for WSC02 and JM#21, although the JM#21/CXCR4 complex is more favored than WSC02/CXCR4 by 10 kcal/mol (Figure S7), displaying an increased contribution of R6 of JM#21 to the binding energy with respect to K6 in WSC02. Similar to the results with EPI-X4, the mutagenesis experiments indicated an improved binding affinity upon introducing the D262A mutation in both cases, WSC02/CXCR4 and JM#21/CXCR4. In the absence of secondary interactions reinforcing the binding of K6/R6 to D262 (unlike the case of D97 and D187), this effect could be related to conformational changes in the mutated receptor/peptide complex allowing K6/R6 to establish new salt bridges with other negatively charged residues within the binding pocket. Thus, the increased receptor binding activity of WSC02 and JM#21 as compared to EPI-X4 appears to be due to the optimal positioning of K6/R6 within the binding pocket of CXCR4. In the case of JM#21, the superior activity can be attributed to the R6 group.

**Figure 7.**
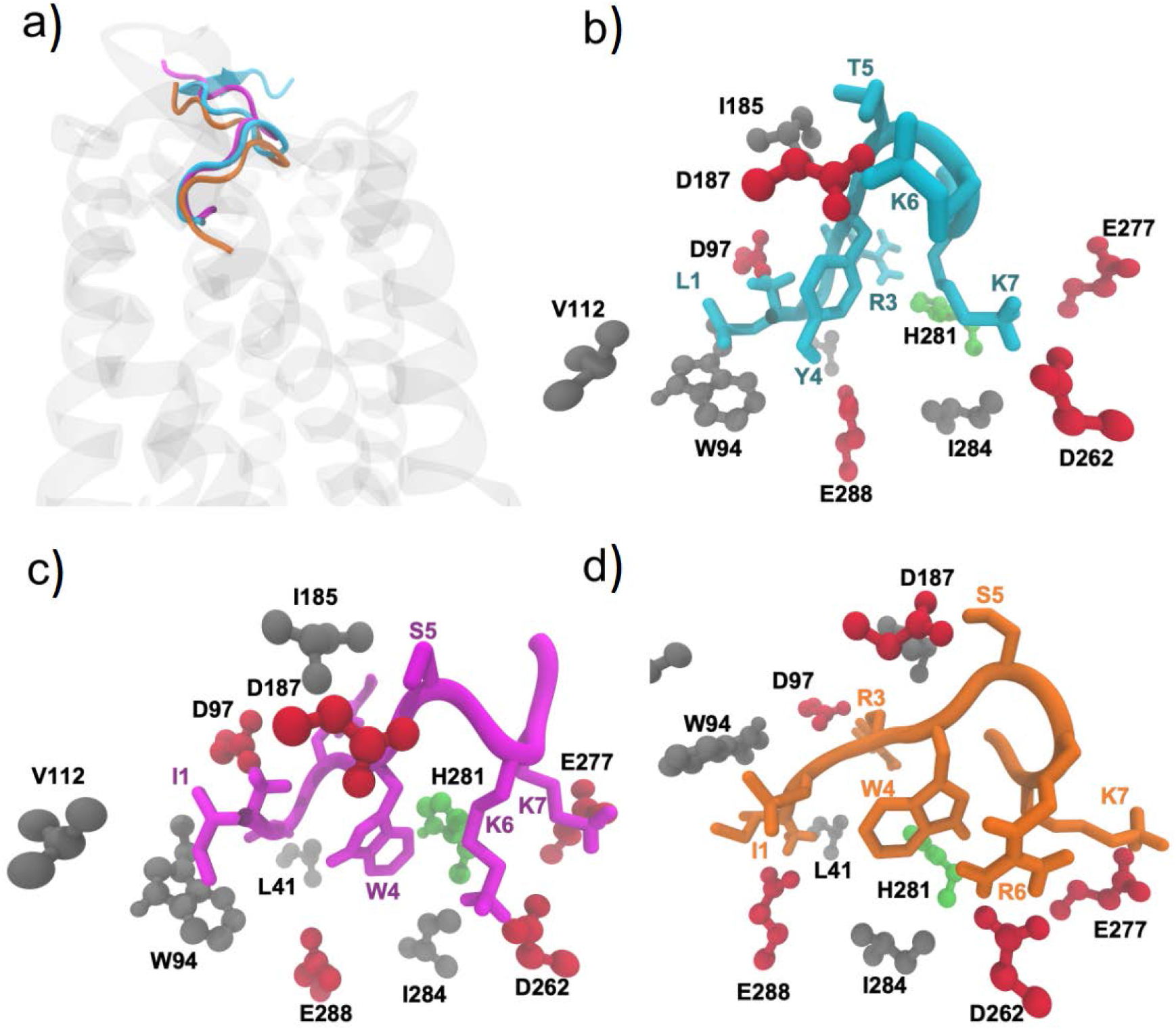
Comparison of the binding modes of EPI-X4 (cyan), WSC02 (purple) and JM#21 (orange). a) The position of the three peptides in the binding pocket, b) EPI-X4, c) WSC02 and d) JM#21 (CPK representation of CXCR4 residues by type: acidic residues = red, polar uncharged = green, nonpolar residues = grey).

### Rational design of shortened EPI-X4 derivatives

As discussed above, our results indicate that only the first 7 amino acids of EPI-X4 and its improved derivatives WSC02 and JM#21 are involved in receptor binding. Thus, truncations as far as up to position 8 might be even possible without a considerable decrease in affinity.

Accordingly, based on the structural knowledge of the binding of EPI-X4 and derivatives to CXCR4 and the individual contributions of specific residues to the binding, we designed a series of C-terminally truncated derivatives of EPI-X4, WSC02 and JM#21 and experimentally assessed their activity (Table 3). Derivatives were truncated by seven or nine residues (EPI-X4), or by three or five residues (WSC02 and JM#21), thereby creating analogues that are nine or seven amino acids long, respectively. In addition, to counteract the electrostatic repulsion of the negatively charged binding pocket of CXCR4, and thus further improve receptor interaction, the negative charge at the C-terminus of the peptides was neutralized by amidation.

**Table 3.** Length optimized derivatives of EPI-X4, WSC02 and JM#21

| Derivative | Sequence | MW | Length | 12G5-competition<br>(nM) ± SEM |
| --- | --- | --- | --- | --- |
| **EPI-X4 (Alb408-423)** | **LVR YTKKVPQVSTPTL** | **1832** | **16** | **1436.4 ± 599.5** |
| EPI-X4 408-416-NH<sub>2</sub> | LVR YTKKVP-NH<sub>2</sub> | 1102 | 9 | 620.6 ± 260.8 |
| EPI-X4 408-414-NH<sub>2</sub> | LVR YTKK-NH<sub>2</sub> | 906 | 7 | 572.6 ± 236.5 |
| **WSC02** | **IVRWSKKVPCVS** | **1401** | **12** | **272.5 ± 147.3** |
| WSC02 408-416-NH<sub>2</sub> | IVRWSKKVP-NH<sub>2</sub> | 1111 | 9 | 220.7 ± 121.7 |
| WSC02 408-414-NH<sub>2</sub> | IVRWSKK-NH<sub>2</sub> | 915 | 7 | 205.2 ± 117.0 |
| **JM#21** | **ILRWSRKLP CVS** | **1458** | **12** | **144.8 ± 19.8** |
| JM#21 408-416-NH<sub>2</sub> | ILRWSRKLP-NH<sub>2</sub> | 1167 | 9 | 262.4 ± 64.9 |
| JM#21 408-414-NH<sub>2</sub> | ILRWSRK-NH<sub>2</sub> | 957 | 7 | 156.8 ± 26.0 |

These peptides with serially truncated C-terminus were tested again for CXCR4 receptor affinity using the antibody competition assay.^32^ Interestingly, C-terminal truncation of EPI-X4 of up to nine amino acids (EPI-X4 408-414) did not lead to a decreased but rather increased binding affinity to CXCR4 (Figure S8). As expected, elimination of K7 (EPI-X4 408-413), almost completely abolished receptor binding, confirming our computational modeling results. Taken together, this suggests that the N-terminal segment of EPI-X4 is highly conserved for the recognition of CXCR4, in agreement with our results indicating that NTER-IN is the most favored binding mode.

Truncated versions of EPI-X4 and WSC02 replaced the CXCR4 antibody 12G5 with a slightly increased activity. For JM#21, the shorter 7 amino acid long analogue interacted with the receptor as strong as the 12 amino acids peptide (Figure 8, Table 3). Although solution structures cannot be directly extrapolated to the situation in which the peptide is bound to CXCR4, the reason for this difference might be found by looking at the conformational flexibility of the peptides. In the solution structures of EPI-X4^16^ and WSC02^17^ the C-terminus and other residues interact with the N-terminus (Leu1, Ile1) within the peptide. In contrast, the NMR structure of JM#21 reveals a free and flexible N-terminus, which might be more available for receptor binding. With truncation of the C-terminus of EPI-X4 and WSC02, the peptide intramolecular H-bonds engaging the N-terminus are interrupted. This modification may make their respective N-termini more flexible thereby ease receptor interaction. Since the N-terminus was already available in JM#21, this effect is inconsequential for this peptide, with no further increase of affinity upon truncation.

**Figure 8.**
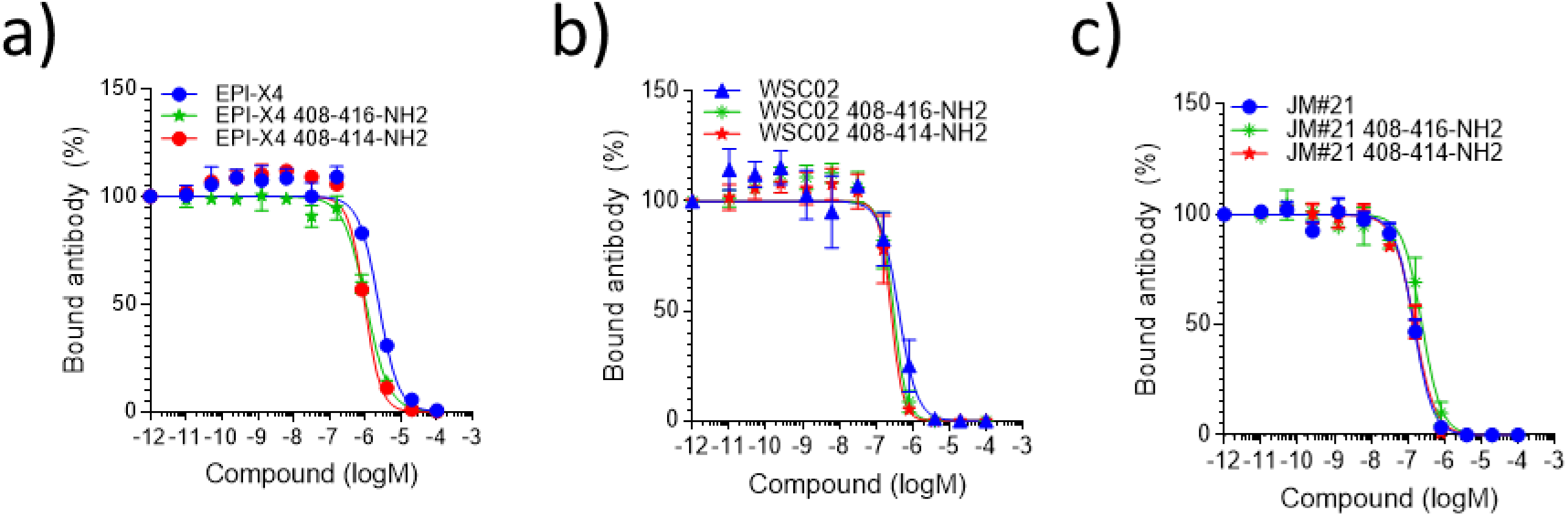
Rationally designed C-terminally truncated EPI-X4 peptides compete with 12G5 antibody binding to CXCR4. EPI-X4 (a), WSC02 (b), or JM#21 (c) and truncated versions thereof were serially diluted and added to SupT1 cells together with a constant concentration of CXCR4 antibody (clone 12G5). After 2 hours, the unbound antibody was removed, and the remaining antibody analyzed in flow cytometry. Shown are data derived from 3 or 4 (JM#21 and derivatives) individual experiments ± SEM.

Next, we tested if those length optimized derivatives are also functionally active CXCR4 antagonists. Downstream signaling of CXCR4 is activated by CXCL12 binding and involves phosphorylation of the signaling proteins Erk and Akt. In the presence of JM#21 and its truncated derivatives CXCL12-mediated activation of both signaling proteins is dose dependently blocked (Figure 9). To see, if also chemotaxis can be effectively inhibited, we incubated cancer T lymphoblasts with different concentrations of JM#21 and its truncated analogues and tested for migration towards physiological concentrations of CXCL12 (Figure 10). As expected, JM#21 inhibited cell migration in a dose-dependent manner, showing 90 ± 6 % inhibition at a concentration of 10 μM, in agreement with previous results.^17^ Interestingly, the shorter versions of JM#21 (408-416-NH2 and 408-414-NH2) inhibited migration almost completely at the same concentration (96 ± 2 % and 99 ± 1 %, respectively) and already to 55 ± 6 and 88 ± 2 % at a concentration of 1 μM.

**Figure 9.**
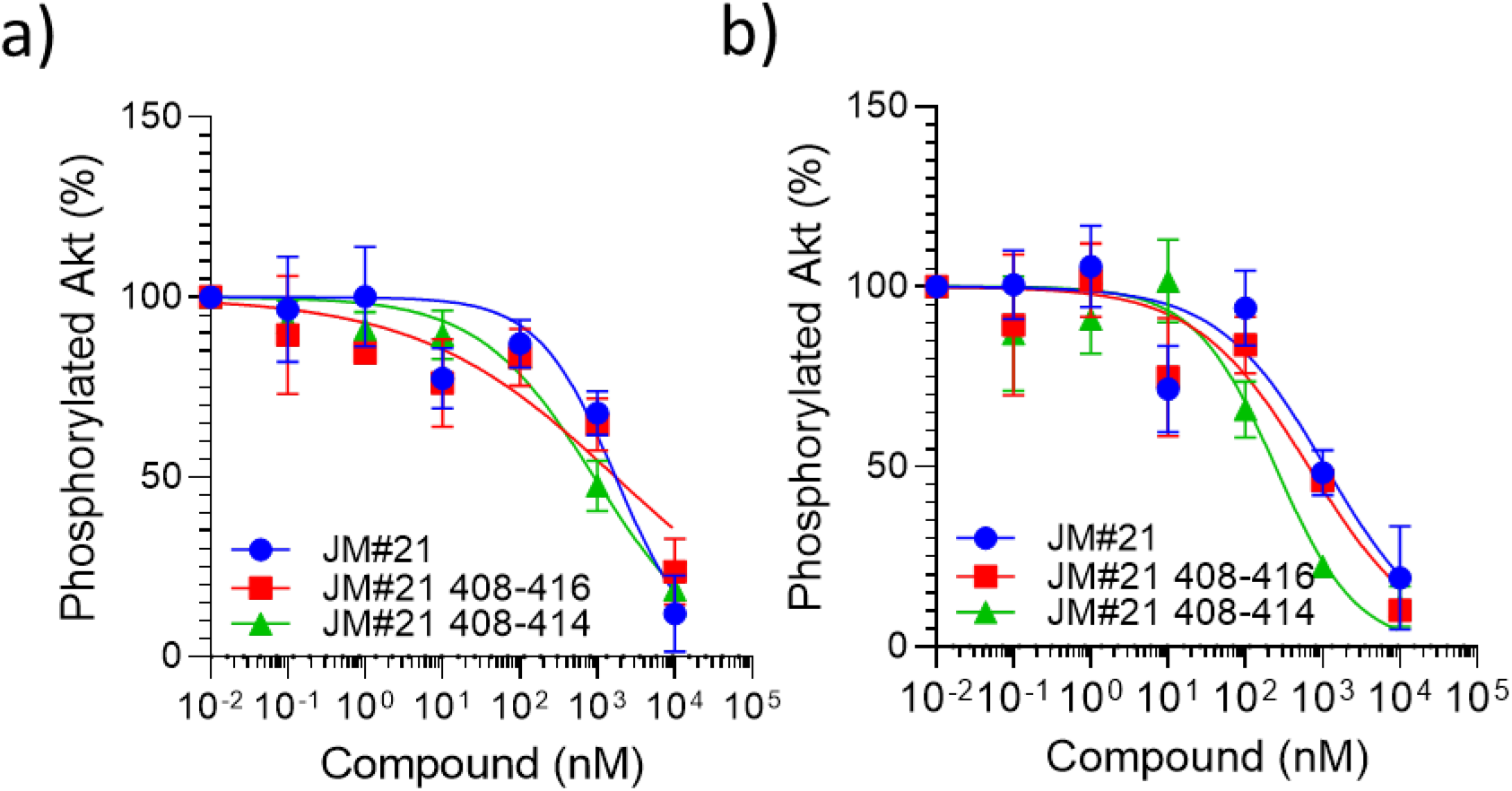
Truncated JM#21 variants does-dependently inhibit CXCL12-mediated signaling. SupT1 cells were stimulated with 100 ng/ml CXCL12 in the presence of peptides for 2 min. Afterwards reaction was stopped by adding 2% PFA and shifting the cells to 4°C. Cells were then permeabilized and subsequently stained with antibodies against pAkt (a) and pErk (b) for analysis in flow cytometry. Shown are data derived from 3 individual experiments ± SEM.

**Figure 10.**
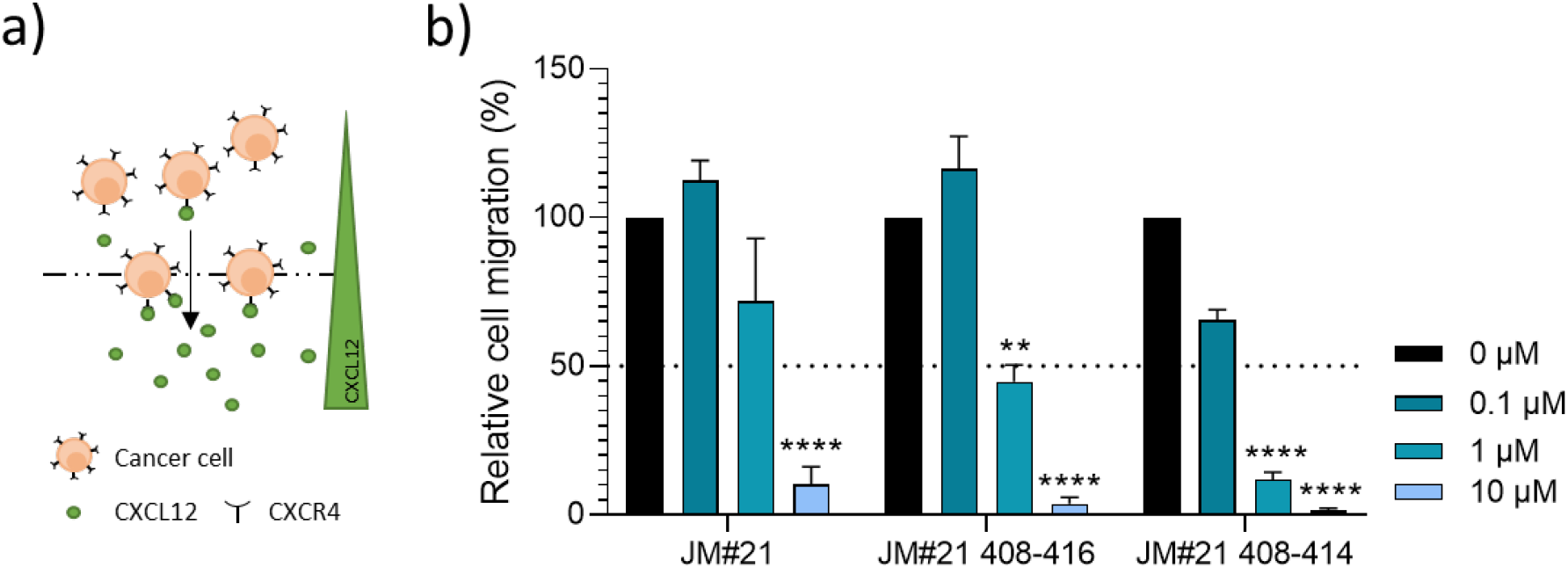
Truncated EPI-X4 JM#21 variants dose-dependently inhibit CXCL12 induced migration of cancer T cells. a) The migration of SupT1 cells towards a 100 ng/ml CXCL12 gradient in a transwell was tested in the presence of indicated concentrations of peptides. After 4 hours amounts of migrated cells were determined by CellTiterGlo ^®^ assay and normalized to values obtained for the PBS control. Shown are values derived from 3 individual experiments performed in triplicates ± SEM. **** p < 0.0001, ** p < 0.01 (one-way ANOVA with Dunnett’s multiple comparison test, compared to PBS control).

Next, we evaluated the toxicity of JM#21 408-414-NH2. To this end, we used zebrafish embryos, which provide a useful *in vivo* model for evaluating toxicity. Zebrafish embryos were exposed to the peptide for 24 hrs starting at 24 hrs post fertilization (hpf), when most organ systems have already developed and are functional. The transparency of the embryos allowed evaluating not only the mortality, but also the sublethal toxicity causing necrosis or lysis (acute toxicity / cytotoxicity), heart edema or reduced / absent circulation (cardiotoxicity), developmental delay or malformations (developmental toxicity) or reduced / absent touch escape response (neurotoxicity) under a light microscope. A standardized scoring system (see experimental details) together with the possibility of investigating embryos on a large scale, yields statistically solid and reproducible results. No toxic effects were found at concentrations that are active in other assays (Figure 11).

**Figure 11.**
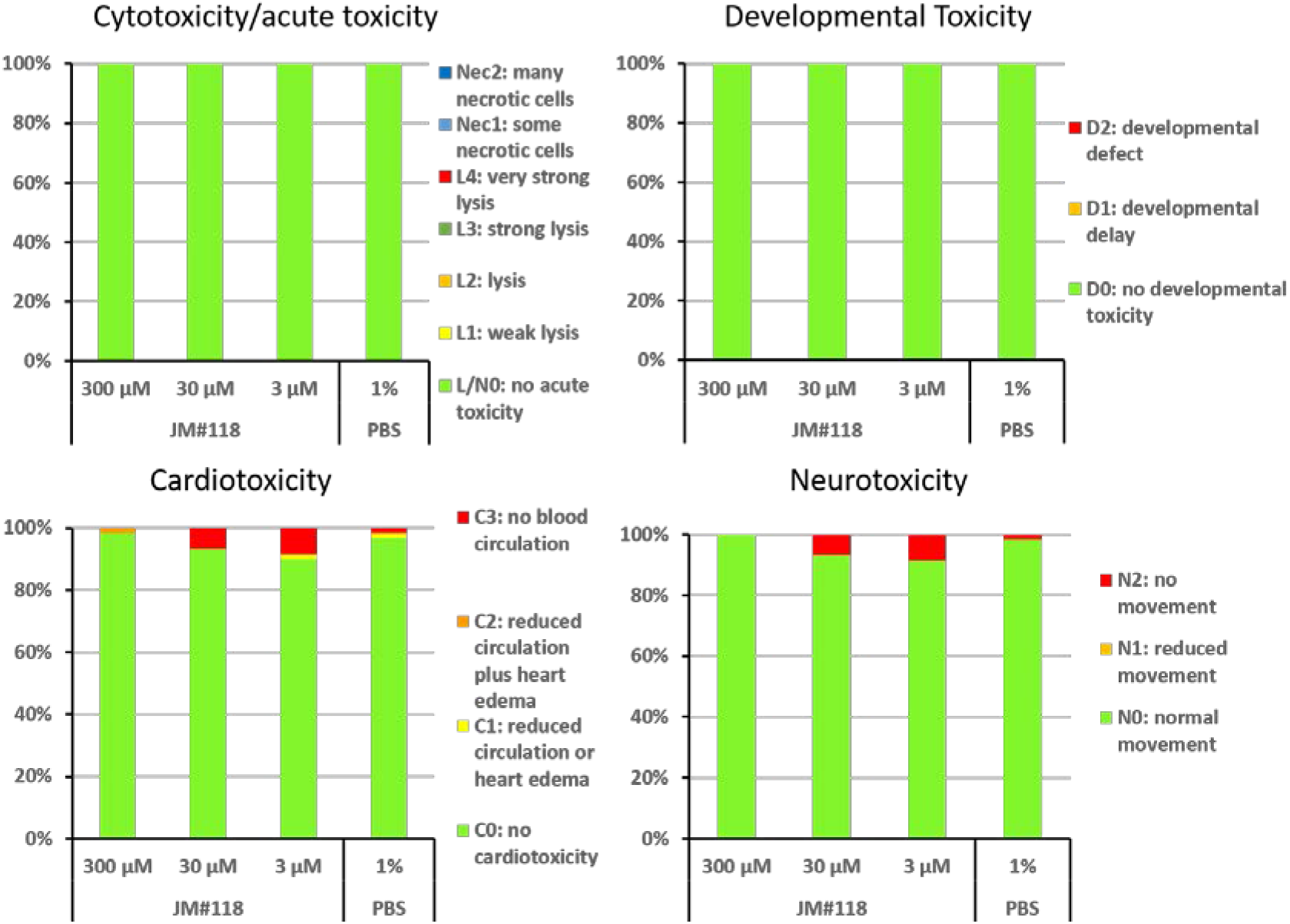
JM#21 408-414-NH2 (JM#118) is not toxic to zebrafish embryos. Zebrafish embryos were scored for mortality or altered phenotypes at 48 hpf after exposure for 24 hrs to the peptide or the negative control (PBS) at the indicated concentrations. Altered phenotypes include necrosis and non-lethal lysis (cytotoxicity), heart edema, reduced or absent circulation (cardiotoxicity), delayed development or malformations (developmental toxicity) and reduced or absent touch escape response (neurotoxicity). n= 60 embryos each group.

Thus, our novel truncated forms of JM#21 are more potent in terms of inhibiting CXCR4-downstream signaling and cancer cell migration, while no toxic effects could be observed as shown for the shortest JM#21 derivative in the zebrafish toxicity assay.

## CONCLUSION

We here present an experimentally proven structural model of the interaction of endogenous EPI-X4 with its receptor CXCR4, based on atomistic biomolecular simulations. EPI-X4 is the first endogenous peptide antagonist of a GPCR that likely plays a key role in CXCR4 regulation. Our results show that EPI-X4 and its improved derivatives interact with CXCR4 mainly via the first seven amino-acid residues, forming interactions with the minor binding pocket of the receptor, in a manner somewhat similar to the binding of viral chemokine antagonist vMIP-II (PDB ID: 4RWS)^11^. These interactions not only occupy the receptor in way that prevents 12G5 antibody binding to ECL2 but also binding of CXCL12, which explains EPI-X4’s CXCR4 antagonizing activity.

Inhibition of CXCR4 is a promising strategy for the treatment of several disorders such as cancers, HIV, and inflammatory diseases. Here, we rationally designed shortened and highly active analogs of EPI-X4 JM#21, a previous lead compound that showed potent therapeutic effects upon topical application in animal models of CXCR4 associated inflammatory diseases, i.e. atopic dermatitis and allergic asthma^17^. The truncated EPI-X4 JM#21 derivatives antagonized cancer cell migration towards CXCL12 even more potently than the precursor peptide. The shortest and most active version (JM#21 408-414-NH2), which exhibited no toxicity in zebrafish assays, encompasses only 7 amino acids and has a molecular weight of 957 Da. Interestingly, almost no orally administered drugs and clinical candidates exist with molecular weights exceeding 1000 Da^49^. Thus, our rationally designed truncated EPI-X4 derivatives should be easier to handle, with less production costs but may also pave the way for oral administration of this new class of CXCR4 antagonists.

## COMPUTATIONAL AND EXPERIMENTAL SECTION

### Computational details

#### CXCR4 model

The crystal structure of CXCR4 (PDB ID: 3ODU)^12^ was used for the modelling studies. However, the N-terminal loop segment (corresponding to residues 1-26) did not have interpretable electron density and therefore this segment is missing in the crystal structure. Although this segment is not crucial for the interaction of small molecular inhibitors, it might influence the binding of larger molecules such as peptides. For this reason, we modelled the complete structure of the CXCR4 protein by combining the coordinates from the crystal structure with the N-terminal region obtained from NMR studies (PDB ID: 2K04)^29^.

#### Docking of the EPI-X4 peptide

The structure of EPI-X4 is available from NMR studies (PDB ID: 2N0X)^16^ in solution. From the solution structure ensemble of EPI-X4, three randomly chosen conformations were the starting point for the docking calculations with CXCR4. The HADDOCK webserver was used for protein-peptide docking studies.^33^ In all cases, residues located at both the minor and major pockets of CXCR4 were chosen as active residues, which serve as the potential site for the interaction with EPI-X4. In the case of EPI-X4, all the residues were considered as active. Default docking parameters corresponding to Easy Interface in HADDOCK were used.^33^ In addition, one binding mode was built by multi-template homology modelling^34^ using the crystal structure CXCR4 complexed with a viral chemokine antagonist vMIP-II (PDB ID: 4RWS)^11^ and the crystal structure 3ODU^12^ as second template. The Modeller program (version 9.18) was used for the homology modelling.^35^

#### Atomistic molecular dynamics simulations

Four different binding modes of EPI-X4 were used as starting points for atomistic molecular dynamics simulations in the lipid bilayer and water. The system consisted of 257 POPC lipids, ~40000 TIP3P water molecules, 50 mM KCl and the CXCR4/EPI-X4 complex and was generated using the CHARMM-GUI server.^35^ The initial system was subjected to energy minimization steps and equilibration molecular dynamics (MD) simulations prior to the production MD runs. For the initial run, harmonic position restraints with a force constant of 10 kcal/mol/Å^2^ were applied on the protein atoms and the atoms of the lipid head groups. The force constant was gradually reduced to zero in six steps of 200 ps equilibration runs. Production MD simulations were carried out using the equilibrated system. In all cases, periodic boundary conditions were used to eliminate surface effects and the particle mesh Ewald (PME)^36^ method was employed for the computation of long-range electrostatics. Short-range Lennard-Jones and electrostatic interactions were cutoff at 12 Å and a switching function was used between 10 and 12 Å to smoothen the interactions at the cutoff distance. Langevin dynamics was employed with the temperature maintained at 300 K and the Langevin piston Nose-Hoover method was used for maintaining the pressure at 1 atm.^37,38^ A time step of 2 fs was used for the integration of the equations of motion. All the simulations were replicated three times using different initial random velocities. The CHARMM36 force field^39,40^ and the NAMD program (version 2.11)^41^ were used.

#### Coarse-grained MD simulations

The CXCR4 receptor complexed with the EPI-X4 derivative (EPI-X4D: 408-416) was simulated at the coarse-grained (CG) level in a POPC lipid-water environment. The CHARMM-GUI server was used to generate the initial configurations.^42^ We employed the Martini 2.2 CG force field^30^ implemented for GROMACS program (version 2016.3)^43^. To reduce the system size, the N-terminal loop (corresponding to residues 1-32) and the C-terminal loop (residues 304-319) were removed and the termini were set as neutral in the CG model. The truncated CXCR4 along with the EPI-X4 derivative (EPI-X4D) were embedded in a POPC bilayer (consisting of 254 lipids) and solvated with an uncharged water model.^30^ The total charge of the system was neutralized by the addition of nine chloride ions. In total, the system contained ~10000 CG particles. Constraints were applied to keep the regular secondary structures intact. NPT simulations were performed using velocity rescale thermostat at 310 K and a Berendsen barostat at 1 atm.^44,45^ The relative dielectric constant, ε_r_=15 was used to account for the screening effect of uncharged water. The short-range LJ interactions were cutoff at 11 Å and shift scheme was used to smoothen the potential. The electrostatic interactions were computed with the reaction-field approach^46^, using a cutoff of 11 Å. A time step of 20 fs was used for the integration of position and velocities.

Three equilibration MD simulations were performed with position restraints on the secondary structures using force constants of 10, 5 and 1 kcal/mol/Å^2^, respectively. The production MD was performed with position restraints operating only on the transmembrane helices of CXCR4 with a force constant of 0.1 kcal/mol/Å^2^. This was necessary to restrict the excessive displacement of the helices.

#### Pulling simulations

To identify the possible binding modes of EPI-X4D, the peptide was pulled from the solution phase to the binding pockets by constant-force MD simulations at the CG level. For this purpose, we defined a reference point located between the major and minor binding pockets (obtained as the center of geometry of four binding pocket residues D187, E288, D262 and D97). The coordinates of this point served as the absolute reference for the pulling simulations. The peptides were kept in the solution phase and at a distance of ~30 Å from the reference point. A 1 μs MD simulation was performed applying the position restraints on the central residue of the peptide (kf=10 kcal/mol/Å^2^) as well as on the TM-helices (k_f_=0.1 kcal/mol/Å^2^). From this simulation, 20 snapshots (with the interval of 50 ns) were taken for the pulling simulations.

Different force constants ranging from 0.025 to 2.5 kcal/mol/Å^2^ were tested for constant force simulations. Constant force simulations were performed with the pulling force operating only on the z-direction. Different points belonging to the peptide (N-terminal residue, C-terminal residue and central residue) were used in different simulations (see Results and Discussion).

### Experimental details

#### Site-directed CXCR4 mutagenesis and cloning experiments

The human CXCR4 gene (isoform 1 or b, NCBI Reference Sequence: NP_003458.1, UniProtKB/Swiss-Prot: P61073-1) was amplified by PCR of the pTrip_GFP_CXCR4 vector (kindly provided by Prof. Françoise Bachelerie, Paris, France) by generating the flanking single cutter sites NheI and HindIII. The PCR-fragment was ligated in the empty pcDNA3.1(+) vector (Life Technologies GmbH, Darmstadt). Afterward the IRES-eGFP cassette of the proviral clone pBR_NL4-3_IRES-eGFP^47^ was PCR-amplified with EcoRI and NotI single cutter sites and ligated in the multiple cloning site after CXCR4. Site-directed mutagenesis (New England Biolabs, E0554S) was used to introduce different point mutations in this construct (L41A, D97E, D97S, D97T, D97N, D97Q, V112A, V112L, D187E, D262A, E277A). Additionally, for some constructs a second and third round of site-directed mutagenesis was performed to introduce two or three amino acid changes in CXCR4 (D97E plus V112A, D97E plus V112L, L41A plus V112A, L41A plus D97E plus V112A). As a negative control, a CXCR4 construct harboring two stop codons after the start codon and a mutation introducing a frameshift was cloned (vector only control). All primers are listed in Table S5. All constructs were sequenced to verify their accuracy.

#### Peptide synthesis

Peptides were synthesized on a 0.10 mM scale using standard Fmoc solid-phase peptide synthesis techniques with the microwave synthesizer (liberty blue; CEM). Peptides were purified using reverse phase preparative high-performance liquid chromatography (HPLC). Peptides were lyophilized, mass was verified by liquid chromatography mass spectroscopy (LCMS) and peptide resolved in PBS before usage.

#### Cell culture

HEK293T cells were cultured in DMEM supplemented with 10 % fetal calf serum (FCS), 100 units/ml penicillin, 100 μg/ml streptomycin, and 2 mM L-glutamine (Gibco). SupT1 suspension cells were cultured in RPMI supplemented with 10 % FCS, 100 units/ml penicillin, 100 μg/ml streptomycin, 2 mM L-glutamine and 1 mM HEPES (Gibco).

#### Antibody competition assay

To test for peptide interaction with CXCR4, 50,000 SupT1 cells were seeded in 96-well microtiter plates in PBS supplemented with 1% FCS, buffer was removed and cells were precooled at 4°C. Peptides were diluted in precooled PBS before 15 μl were added to the cells together with 15 μl APC-conjugated CXCR4 antibody (clone 12G5; #555976, PD PharmingenTM) at a concentration close to its EC_50_. Cells were incubated at 4°C for 2 hours. Afterwards, unbound antibody was removed and cells analyzed by flow cytometry. The mean fluorescence intensity (MFI) was normalized to the PBS control. Values were normalized to PBS control + antibody. IC_50_ values were determined by non-linear regression using GraphPad Prism. Antibody competition with CXCR4 mutants 293T cells were transiently transfected with pcDNA3.1 containing eGFP and CXCR4 wt or CXCR4 harboring selected point mutations. As control the vector without functional CXCR4 (vector only) or only transfection reagent (mock) was used. The next day, medium was changed. Cells were harvested one day after, washed and 50,000 cells seeded in V-well microtiter plates. Antibody competition was performed as described before.^32^ For analysis after flow cytometry eGFP expressing cells were gated and further analyzed. CXCR4 expression levels and antibody binding to CXCR4 was determined using the CXCR4 antibody clones 12G5 (binds to the second extracellular loop) (#555976, BD PharmingenTM) and 1D9 (binds to the N-terminus) (#551510, BD PharmingenTM) (Supplement Figure S3). For the binding experiment, cells with impaired antibody binding or eGFP expression levels compared to the wild type control were excluded. After antibody competition, MFIs were normalized to the PBS control + antibody. IC_50_ values were determined by non-linear regression. **** p < 0.0001, *** p < 0.001, ** p < 0.01, * p < 0.1 (one-way ANOVA with Dunnett’s multiple comparison test).

#### ERK/AKT signaling assay

CXCL12 induced ERK and AKT phosphorylation was determined in SupT1 cells. For this, 100,000 cells were seeded per well in a 96-V well plate in 100 μl medium supplemented with 1% FCS. Cells were incubated for 2 hours at 37°C before 5 μl of compounds were added. After 15 min incubation at 37°C cells were stimulated by adding 5 μl CXCL12 diluted in PBS to reach a final concentration of 100 ng/ml. Cells were further incubated for 2 min before the reaction was stopped by adding 20 μl of 10% PFA. Cells were fixed for 15 min at 4°C before PFA was removed and cell permeabilized by adding 100 μl ice cold methanol. After 15 min at 4°C the methanol was removed, cells were washed and 30 μl primary antibody was added (phospho-p44/42 MAPK (Erk1) (Tyr204)/ (Erk2) (Tyr187) (D1H6G) mouse mAb #5726; phosphor-Akt (Ser473) (193H12) rabbit mAb #4058 Cell Signaling) for 1 hour at 4°C. After the antibody was removed and cells were washed secondary antibody was added for 30 min. Cells were washed afterwards and subsequently analyzed by flow cytometry.

#### Migration of cancer T lymphoma cells

Migration assays towards a 100 ng/ml CXCL12 gradient (#300-28A, Peprotech) were performed using 96-well transwell assay plates (Corning Incorporated, Kennebunk, ME, USA) with 5 μm polycarbonate filters. First, 50 μl (0.75 x 105) SupT1 cells resuspended in assay buffer (RPMI supplemented with 0.1 % BSA) were seeded into the upper chamber in the presence or absence of compounds and allowed to settle down for around 15 min. In the meantime, 200 μl assay buffer supplemented with or without 100 ng/ml CXCL12 as well as compounds were filled into a 96 well-V plate. Cells were allowed to migrate towards CXCL12 by putting upper chamber onto the 96 well-V plate. After a migration time of 4 h at 37°C (5 % CO2) the lower compartments were analyzed for cell content by Cell-Titer-Glo^®^ assay (Promega, Madison, WI, USA). Percentages of migrated cells were calculated as described before^48^ and normalized to the CXCL12-only control.

#### Toxicity assays in zebrafish

Wild-type zebrafish embryos were dechorionated at 24 hrs post fertilization (hpf) using digestion with 1 mg/ml pronase (Sigma) in E3 medium (83 μM NaCl, 2.8 μM KCl, 5.5 μM CaCl2, 5.5 μM MgSO4). Embryos were exposed for 24 hrs, in groups of 3, to 100 μl of E3 containing JM#21 408-414-NH2 at 3, 30 and 300 μM. Each concentration was tested in two independent assays, each of which was performed on 10 × 3 embryos. The peptide solvent (PBS), diluted in E3, was used as negative control at the same amount as introduced by the highest peptide concentration. As positive control for acute toxicity/cytotoxicity the pleurocidin antimicrobial peptide NRC-03 (GRRKRKWLRRIGKGVKIIGGAALDHL-NH2) was used at a concentration of 6 μM as described^50^. Abamectin at a concentration of 3.125 μM was used as positive control for neurotoxicity^51^. At 48 hpf (after 24 hrs of incubation) embryos were scored in a stereomicroscope for signs of acute toxicity/cytotoxicity (lysis and/or necrosis), developmental toxicity (delay and/or malformations), or cardiotoxicity (heart edema and/or reduced or absent circulation). Each embryo was also touched with a needle and reduced or absent touch response (escape movements) was evaluated as signs of neurotoxicity if and only if no signs of acute toxicity were present in the same embryo. Embryos were categorized within each of these toxicity categories into several classes of severity according to the criteria listed in Table S6. The Chi-Square test was used to calculate whether the distribution of embryos into toxicity classes differed significantly between the PBS negative control and the test substances.

## ASSOCIATED CONTENT

### Supporting information

(i) Molecular dynamics simulation data; RMSD and RMSF analysis, H-bonding data, clustering analysis, and surface area plots corresponding to EPI-X4, WSC02, and JM#21 peptides, (ii) Experimental data for CXCR4 expression, antibody assay, IC_50_ for EPI-X4, WSC02 and JM#21 peptides, (iii) NMR spectroscopy methodology and results for the EPI-X4 peptide and (iv) toxicity categories are available free of charge.

### Author Contributions

P. S. performed the simulations, analyzed the data and wrote the manuscript together with M.H. and E.S-G. M. H. performed the experiments, analyzed the experimental data and wrote the manuscript together with P.S. and E.S-G. C.S. did the cloning experiments, A.G. performed the migration essays, G. K. contributed the NMR studies, N.P and L. S. did the peptide synthesis and purity control. M.R. and G.W. performed the zebrafish toxicity studies. J.M. coordinated and supervised the experimental work and designed the project together with E.S-G. E.S-G designed the project, coordinated and supervised the work, analyzed the data and wrote the manuscript with P.S and M. H.

## ACKNOWLEDGMENT

This work was supported by the Deutsche Forschungsgemeinschaft (DFG, German Research Foundation) under the collaborative research center CRC 1279 (project A06). E. S.-G. was also supported by Germany’s Excellence Strategy – EXC 2033 – 390677874 – RESOLV and the collaborative research center CRC 1093 “Supramolecular Chemistry on Proteins”, both funded by the DFG. E. S.-G. acknowledges the support of the Boehringer Ingelheim Foundation (Plus-3 Program) and the computational time provided by the Computing and Data Facility of the Max Planck Society and the supercomputer magnitUDE of the University of Duisburg-Essen. P.S. acknowledges the Department of Science and Technology in India for the support received through DST-SERB SRG/2019/0.02156. J.M. further acknowledges funding by the Baden-Württemberg Stiftung, the European Research Council, and the DFG grant MU 3315/11-1. M.H. is part of the International Graduate School in Molecular Medicine Ulm.

## ABBREVIATIONS

CXCR4: C-X-C chemokine receptor 4
MD: molecular dynamics
CG: coarse-grained
CXCR4: antagonist

## REFERENCES

(1) Pozzobon, T.; Goldoni, G.; Viola, A.; Molon, B. CXCR4 Signaling in Health and Disease. Immunol. Lett. 2016, 177, 6–15. https://doi.org/10.1016/j.imlet.2016.06.006.

(2) Garc\’\ia-Cuesta, E. M.; Santiago, C. A.; Vallejo, J.; Juarranz, Y.; Rodriguez Frade, J. M.; Mellado, M. The Role of the CXCL12/CXCR4/ACKR3 Axis in Autoimmune Diseases. Front. Endocrinol. (Lausanne). 2019, 10, 585.

(3) Choi, W.-T.; Duggineni, S.; Xu, Y.; Huang, Z.; An, J. Drug Discovery Research Targeting the CXC Chemokine Receptor 4 (CXCR4). J. Med. Chem. 2012, 55 (3), 977–994. https://doi.org/10.1021/jm200568c.

(4) Choi, W.-T.; An, J. Biology and Clinical Relevance of Chemokines and Chemokine Receptors CXCR4 and CCR5 in Human Diseases. Exp. Biol. Med. 2011, 236 (6), 637–647.

(5) Chatterjee, S.; Behnam Azad, B.; Nimmagadda, S. Chapter Two - The Intricate Role of CXCR4 in Cancer. In Emerging Applications of Molecular Imaging to Oncology; Pomper, M. G., Fisher, P. B., Eds.; Advances in Cancer Research; Academic Press, 2014; Vol. 124, pp 31–82. https://doi.org/10.1016/B978-0-12-411638-2.00002-1.

(6) Zhang, J.; Liu, C.; Mo, X.; Shi, H.; Li, S. Mechanisms by Which CXCR4/CXCL12 Cause Metastatic Behavior in Pancreatic Cancer. Oncol Lett 2018, 15 (2), 1771–1776. https://doi.org/10.3892/ol.2017.7512.

(7) Mukherjee, D.; Zhao, J. The Role of Chemokine Receptor CXCR4 in Breast Cancer Metastasis. Am. J. Cancer Res. 2013, 3 (1), 46.

(8) Pawig, L.; Klasen, C.; Weber, C.; Bernhagen, J.; Noels, H. Diversity and Inter-Connections in the CXCR4 Chemokine Receptor/Ligand Family: Molecular Perspectives. Front. Immunol. 2015, 6, 429. https://doi.org/10.3389/fimmu.2015.00429.

(9) Bleul, C. C.; Wu, L.; Hoxie, J. A.; Springer, T. A.; Mackay, C. R. The HIV Coreceptors CXCR4 and CCR5 Are Differentially Expressed and Regulated on Human T Lymphocytes. Proc. Natl. Acad. Sci. 1997, 94 (5), 1925–1930.

(10) Hendrix, C. W.; Collier, A. C.; Lederman, M. M.; Schols, D.; Pollard, R. B.; Brown, S.; Jackson, J. B.; Coombs, R. W.; Glesby, M. J.; Flexner, C. W.; others. Safety, Pharmacokinetics, and Antiviral Activity of AMD3100, a Selective CXCR4 Receptor Inhibitor, in HIV-1 Infection. JAIDS J. Acquir. Immune Defic. Syndr. 2004, 37 (2), 1253–1262.

(11) Qin, L.; Kufareva, I.; Holden, L. G.; Wang, C.; Zheng, Y.; Zhao, C.; Fenalti, G.; Wu, H.; Han, G. W.; Cherezov, V.; others. Crystal Structure of the Chemokine Receptor CXCR4 in Complex with a Viral Chemokine. Science (80-.). 2015, 347 (6226), 1117–1122.

(12) Wu, B.; Chien, E. Y. T.; Mol, C. D.; Fenalti, G.; Liu, W.; Katritch, V.; Abagyan, R.; Brooun, A.; Wells, P.; Bi, F. C.; others. Structures of the CXCR4 Chemokine GPCR with Small-Molecule and Cyclic Peptide Antagonists. Science (80-.). 2010, 330 (6007), 1066–1071.

(13) Roumen, L.; Scholten, D. J.; de Kruijf, P.; de Esch, I. J. P.; Leurs, R.; de Graaf, C. C(X)CR in Silico: Computer-Aided Prediction of Chemokine Receptor–Ligand Interactions. Drug Discov. Today Technol. 2012, 9 (4), e281–e291. https://doi.org/10.1016/j.ddtec.2012.05.002.

(14) Stephens, B. S.; Ngo, T.; Kufareva, I.; Handel, T. M. Functional Anatomy of the Full Length CXCR4-CXCL12 Complex Systematically Dissected by Quantitative Model-Guided Mutagenesis. BioRxiv 2020.

(15) Buske, C.; Kirchhoff, F.; Münch, J. EPI-X4, a Novel Endogenous Antagonist of CXCR4. Oncotarget 2015, 6 (34), 35137.

(16) Zirafi, O.; Kim, K.-A.; Ständker, L.; Mohr, K. B.; Sauter, D.; Heigele, A.; Kluge, S. F.; Wiercinska, E.; Chudziak, D.; Richter, R.; Moepps, B.; Gierschik, P.; Vas, V.; Geiger, H.; Lamla, M.; Weil, T.; Burster, T.; Zgraja, A.; Daubeuf, F.; Frossard, N.; Hachet-Haas, M.; Heunisch, F.; Reichetzeder, C.; Galzi, J.-L.; Pérez-Castells, J.; Canales-Mayordomo, A.; Jiménez-Barbero, J.; Giménez-Gallego, G.; Schneider, M.; Shorter, J.; Telenti, A.; Hocher, B.; Forssmann, W.-G.; Bonig, H.; Kirchhoff, F.; Münch, J. Discovery and Characterization of an Endogenous CXCR4 Antagonist. Cell Rep. 2015, 11 (5), 737–747. https://doi.org/10.1016/j.celrep.2015.03.061.

(17) Harms, M.; Habib, M. M. W.; Nemska, S.; Nicolò, A.; Gilg, A.; Preising, N.; Sokkar, P.; Carmignani, S.; Raasholm, M.; Weidinger, G.; Kizilsavas, G.; Wagner, M.; Ständker, L.; Abadi, A.; Jumaa, H.; Kirchhoff, F.; Frossard, N.; Sanchez-Garcia, E.; Münch, J. An Optimized Derivative of an Endogenous CXCR4 Antagonist Prevents Atopic Dermatitis and Airway Inflammation. bioRxiv 2020. https://doi.org/10.1101/2020.08.28.272781.

(18) Trent, J. O.; Wang, Z.; Murray, J. L.; Shao, W.; Tamamura, H.; Fujii, N.; Peiper, S. C. Lipid Bilayer Simulations of CXCR4 with Inverse Agonists and Weak Partial Agonists. J. Biol. Chem. 2003, 278 (47), 47136–47144.

(19) Kawatkar, S. P.; Yan, M.; Gevariya, H.; Lim, M. Y.; Eisold, S.; Zhu, X.; Huang, Z.; An, J. Computational Analysis of the Structural Mechanism of Inhibition of Chemokine Receptor CXCR4 by Small Molecule Antagonists. Exp. Biol. Med. 2011, 236 (7), 844–850. https://doi.org/10.1258/ebm.2011.010345.

(20) Cong, X.; Golebiowski, J. Allosteric Na+-Binding Site Modulates CXCR4 Activation. Phys. Chem. Chem. Phys. 2018, 20 (38), 24915–24920.

(21) Mona, C. E.; Besserer-Offroy, É.; Cabana, J.; Lefrançois, M.; Boulais, P. E.; Lefebvre, M.-R.; Leduc, R.; Lavigne, P.; Heveker, N.; Marsault, É.; Escher, E. Structure–Activity Relationship and Signaling of New Chimeric CXCR4 Agonists. J. Med. Chem. 2016, 59 (16), 7512–7524. https://doi.org/10.1021/acs.jmedchem.6b00566.

(22) Huang, X.; Shen, J.; Cui, M.; Shen, L.; Luo, X.; Ling, K.; Pei, G.; Jiang, H.; Chen, K. Molecular Dynamics Simulations on SDF-1α: Binding with CXCR4 Receptor. Biophys. J. 2003, 84(1), 171–184. https://doi.org/10.1016/S0006-3495(03)74840-1.

(23) Yoshikawa, Y.; Kobayashi, K.; Oishi, S.; Fujii, N.; Furuya, T. Molecular Modeling Study of Cyclic Pentapeptide CXCR4 Antagonists: New Insight into CXCR4--FC131 Interactions. Bioorg. Med. Chem. Lett. 2012, 22 (6), 2146–2150.

(24) Mungalpara, J.; Thiele, S.; Eriksen, Ø.; Eksteen, J.; Rosenkilde, M. M.; Våbenø, J. Rational Design of Conformationally Constrained Cyclopentapeptide Antagonists for C-X-C Chemokine Receptor 4 (CXCR4). J. Med. Chem. 2012, 55 (22), 10287–10291. https://doi.org/10.1021/jm300926y.

(25) Oum, Y. H.; Kell, S. A.; Yoon, Y.; Liang, Z.; Burger, P.; Shim, H. Discovery of Novel Aminopiperidinyl Amide CXCR4 Modulators through Virtual Screening and Rational Drug Design. Eur. J. Med. Chem. 2020, 201, 112479. https://doi.org/10.1016/j.ejmech.2020.112479.

(26) Kharche, S. A.; Sengupta, D. Dynamic Protein Interfaces and Conformational Landscapes of Membrane Protein Complexes. Curr. Opin. Struct. Biol. 2020, 61, 191–197. https://doi.org/10.1016/j.sbi.2020.01.001.

(27) Gahbauer, S.; Pluhackova, K.; Böckmann, R. A. Closely Related, yet Unique: Distinct Homo-and Heterodimerization Patterns of G Protein Coupled Chemokine Receptors and Their Fine-Tuning by Cholesterol. PLoS Comput. Biol. 2018, 14 (3), e1006062.

(28) Rodríguez, D.; Gutiérrez-de-Terán, H. Characterization of the Homodimerization Interface and Functional Hotspots of the CXCR4 Chemokine Receptor. Proteins Struct. Funct. Bioinforma. 2012, 80 (8), 1919–1928. https://doi.org/10.1002/prot.24099.

(29) Veldkamp, C. T.; Seibert, C.; Peterson, F. C.; Norberto, B.; Haugner, J. C.; Basnet, H.; Sakmar, T. P.; Volkman, B. F. Structural Basis of CXCR4 Sulfotyrosine Recognition by the Chemokine SDF-1/CXCL12. Sci. Signal. 2008, 1 (37), ra4--ra4.

(30) de Jong, D. H.; Singh, G.; Bennett, W. F. D.; Arnarez, C.; Wassenaar, T. A.; Schäfer, L. V; Periole, X.; Tieleman, D. P.; Marrink, S. J. Improved Parameters for the Martini Coarse-Grained Protein Force Field. J. Chem. Theory Comput. 2013, 9 (1), 687–697.

(31) Humphrey, W.; Dalke, A.; Schulten, K. VMD: Visual Molecular Dynamics. J. Mol. Graph. 1996, 14 (1), 33–38.

(32) Harms, M.; Gilg, A.; Ständker, L.; Beer, A. J.; Mayer, B.; Rasche, V.; Gruber, C. W.; Münch, J. Microtiter Plate-Based Antibody-Competition Assay to Determine Binding Affinities and Plasma/Blood Stability of CXCR4 Ligands. Sci. Rep. 2020, 10 (1). https://doi.org/10.1038/s41598-020-73012-4.

(33) van Zundert, G. C. P.; Rodrigues, J. P. G. L. M.; Trellet, M.; Schmitz, C.; Kastritis, P. L.; Karaca, E.; Melquiond, A. S. J.; van Dijk, M.; de Vries, S. J.; Bonvin, A. M. J. J. The HADDOCK2.2 Web Server: User-Friendly Integrative Modeling of Biomolecular Complexes. J. Mol. Biol. 2016, 428 (4), 720–725. https://doi.org/10.1016/j.jmb.2015.09.014.

(34) Sokkar, P.; Mohandass, S.; Ramachandran, M. Multiple Templates-Based Homology Modeling Enhances Structure Quality of AT1 Receptor: Validation by Molecular Dynamics and Antagonist Docking. J. Mol. Model. 2011, 17 (7). https://doi.org/10.1007/s00894-010-0860-z.

(35) Eswar, N.; Webb, B.; Marti-Renom, M. A.; Madhusudhan, M. S.; Eramian, D.; Shen, M.; Pieper, U.; Sali, A. Comparative Protein Structure Modeling Using Modeller. Curr. Protoc. Bioinforma. 2006, 15 (1), 5–6.

(36) Darden, T.; York, D.; Pedersen, L. Particle Mesh Ewald: An N· Log (N) Method for Ewald Sums in Large Systems. J. Chem. Phys. 1993, 98 (12), 10089–10092.

(37) Martyna, G. J.; Tobias, D. J.; Klein, M. L. Constant Pressure Molecular Dynamics Algorithms. J. Chem. Phys. 1994, 101 (5), 4177–4189.

(38) Feller, S. E.; Zhang, Y.; Pastor, R. W.; Brooks, B. R. Constant Pressure Molecular Dynamics Simulation: The Langevin Piston Method. J. Chem. Phys. 1995, 103 (11), 4613–4621.

(39) Klauda, J. B.; Venable, R. M.; Freites, J. A.; O’Connor, J. W.; Tobias, D. J.; Mondragon-Ramirez, C.; Vorobyov, I.; MacKerell Jr, A. D.; Pastor, R. W. Update of the CHARMM All-Atom Additive Force Field for Lipids: Validation on Six Lipid Types. J. Phys. Chem. B 2010, 114 (23), 7830–7843.

(40) Best, R. B.; Zhu, X.; Shim, J.; Lopes, P. E. M.; Mittal, J.; Feig, M.; MacKerell Jr, A. D. Optimization of the Additive CHARMM All-Atom Protein Force Field Targeting Improved Sampling of the Backbone ϕ, ψ and Side-Chain X1 and X2 Dihedral Angles. J. Chem. Theory Comput. 2012, 8 (9), 3257–3273.

(41) Phillips, J. C.; Braun, R.; Wang, W.; Gumbart, J.; Tajkhorshid, E.; Villa, E.; Chipot, C.; Skeel, R. D.; Kale, L.; Schulten, K. Scalable Molecular Dynamics with NAMD. J. Comput. Chem. 2005, 26 (16), 1781–1802.

(42) Jo, S.; Kim, T.; Iyer, V. G.; Im, W. CHARMM-GUI: A Web-Based Graphical User Interface for CHARMM. J. Comput. Chem. 2008, 29 (11), 1859–1865.

(43) Van Der Spoel, D.; Lindahl, E.; Hess, B.; Groenhof, G.; Mark, A. E.; Berendsen, H. J. C. GROMACS: Fast, Flexible, and Free. J. Comput. Chem. 2005, 26 (16), 1701–1718. https://doi.org/10.1002/jcc.20291.

(44) Bussi, G.; Donadio, D.; Parrinello, M. Canonical Sampling through Velocity Rescaling. J. Chem. Phys. 2007, 126 (1), 14101.

(45) Berendsen, H. J. C.; Postma, J. P. M. van; van Gunsteren, W. F.; DiNola, A.; Haak, J. R. Molecular Dynamics with Coupling to an External Bath. J. Chem. Phys. 1984, 81 (8), 3684–3690.

(46) Tironi, I. G.; Sperb, R.; Smith, P. E.; van Gunsteren, W. F. A Generalized Reaction Field Method for Molecular Dynamics Simulations. J. Chem. Phys. 1995, 102 (13), 5451–5459.

(47) Schindler, M.; Münch, J.; Kutsch, O.; Li, H.; Santiago, M. L.; Bibollet-Ruche, F.; Müller-Trutwin, M. C.; Novembre, F. J.; Peeters, M.; Courgnaud, V.; others. Nef-Mediated Suppression of T Cell Activation Was Lost in a Lentiviral Lineage That Gave Rise to HIV-1. Cell 2006, 125 (6), 1055–1067.

(48) Balabanian, K.; Lagane, B.; Infantino, S.; Chow, K. Y. C.; Harriague, J.; Moepps, B.; Arenzana-Seisdedos, F.; Thelen, M.; Bachelerie, F. The Chemokine SDF-1/CXCL12 Binds to and Signals through the Orphan Receptor RDC1 in T Lymphocytes. J. Biol. Chem. 2005, 280 (42), 35760–35766.

(49) Doak, B. C.; Over, B.; Giordanetto, F.; Kihlberg, J. Oral Druggable Space beyond the Rule of 5: Insights from Drugs and Clinical Candidates. Chemistry & Biology 2014, 21 (9), 1115–1142.

(50) Morash MG, Douglas SE, Robotham A, Ridley CM, Gallant JW, Soanes KH. The zebrafish embryo as a tool for screening and characterizing pleurocidin host-defense peptides as anti-cancer agents. DMM Dis Model Mech 2011, 4:622–633.

(51) Raftery TD, Isales GM, Yozzo KL, Volz DC. High-content screening assay for identification of chemicals impacting spontaneous activity in zebrafish embryos. Environ Sci Technol 2014, 48:804–810.

