## supplement for "Biomolecular models of EPI-X4 binding to CXCR4 allow the rational optimization of peptides with therapeutic potential"

[**Figure S3.** CXCR4 expression levels in transfected 293T cells. Point mutations were introduced into the sequence of CXCR4 in an IRES-GFP expression vector. Cells were transfected and GPF positive cells were then stained by either the CXCR4 antibody 12G5 (ECL2) or 1D9 (N-terminus) and analyzed by flow cytometry (with the exception of mock control, where also GFP-negative cells were analyzed). Results were normalized to CXCR4 wt expression levels transfected in 293T cells. Shown are data derived from 3 individual experiments ± SEM. # cell were excluded from further binding experiments due to low expression levels, ## no GFP positive cells could be detected.	8](#_Toc51930068)

[**Figure S12.** EPI-X4 blue to red: N-terminus to C-terminus a) with 50 mM NaOAc b) previously published EPI-X4 without NaOAc. The first 2 amino acids of the N-terminus are free and flexible, while the C-terminus is engaged in hydrogen bonds between Thr6 and Gln10 sidechains as well as a hydrogen bond between the backbone carbonyl of Thr6 and the amide hydrogen of Val11. Additionally, the sidechains of Ser12 and Leu16 are engaged in hydrogen bonds. Arg3 builds a hydrogen bond with its sidechain guanidino group and the carboxyl end of the peptide, when the N-terminus comes close to the C-terminus.	18](#_Toc51930077)

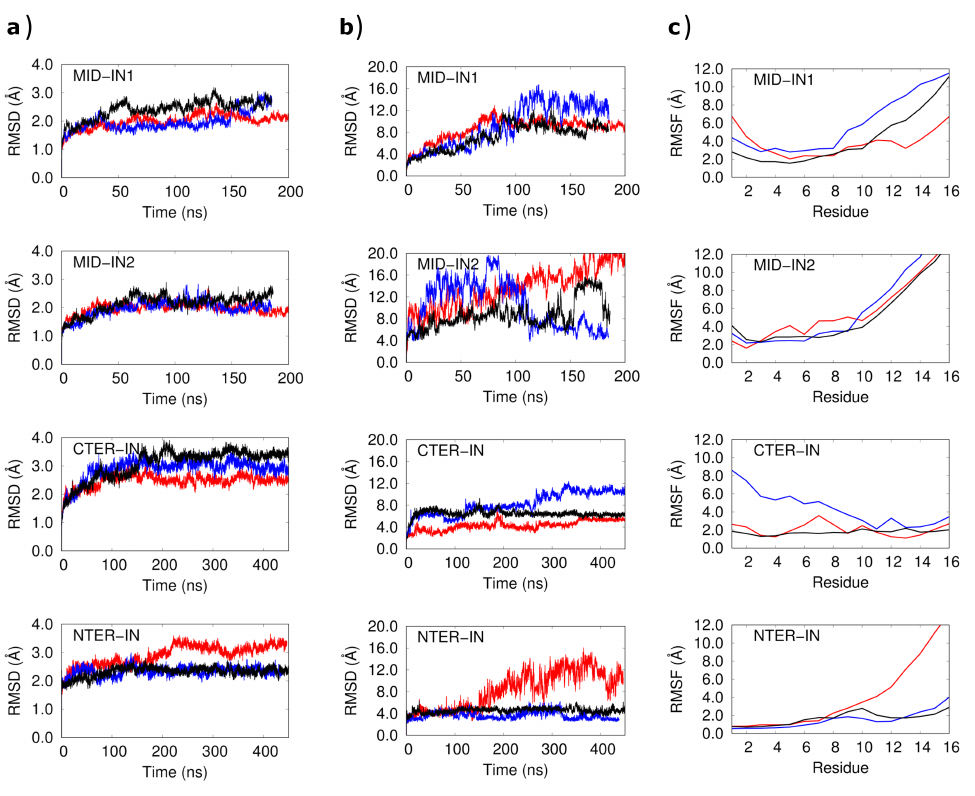

**Figure S1**. Root mean squared deviation (RMSD) and root mean squared fluctuation (RMSF) from the MD simulations of CXCR4/EPI-X4 complexes. a) RMSD of the backbone atoms of CXCR4 during the three replicas of the simulations (red, blue, and black lines). b) RMSD of the backbone atoms of EPI-X4 and c) RMSF of EPI-X4 residues averaged over the simulation time. The analysis was carried out after aligning the backbone atoms of CXCR4 to the initial structure. The N- and C-terminal loops of CXCR4 were omitted in the RMSD analysis.

### Supplementary discussion

**Figure S1:** The analysis of the MDs indicated that the N-terminal and C-terminal loops of CXCR4 are highly disordered, as shown by the root mean squared deviation (RMSD) values. The rest of the protein displayed very low RMSD values (<4 Å), signaling few structural fluctuations (Figure S1a). On the other hand, the RMSDs of EPI-X4 show large variations, especially in case of the MID-IN and MID-IN2 binding modes (Figure S1b). The CTER-IN and NTER-IN conformations exhibited a somewhat stable behavior (Figure S1b), suggesting that these are favored. The root mean squared fluctuation (RMSF) of EPI-X4 evidenced large variations in the C-terminal region in all trajectories corresponding to the MID-IN1 and MID-IN2 binding modes.

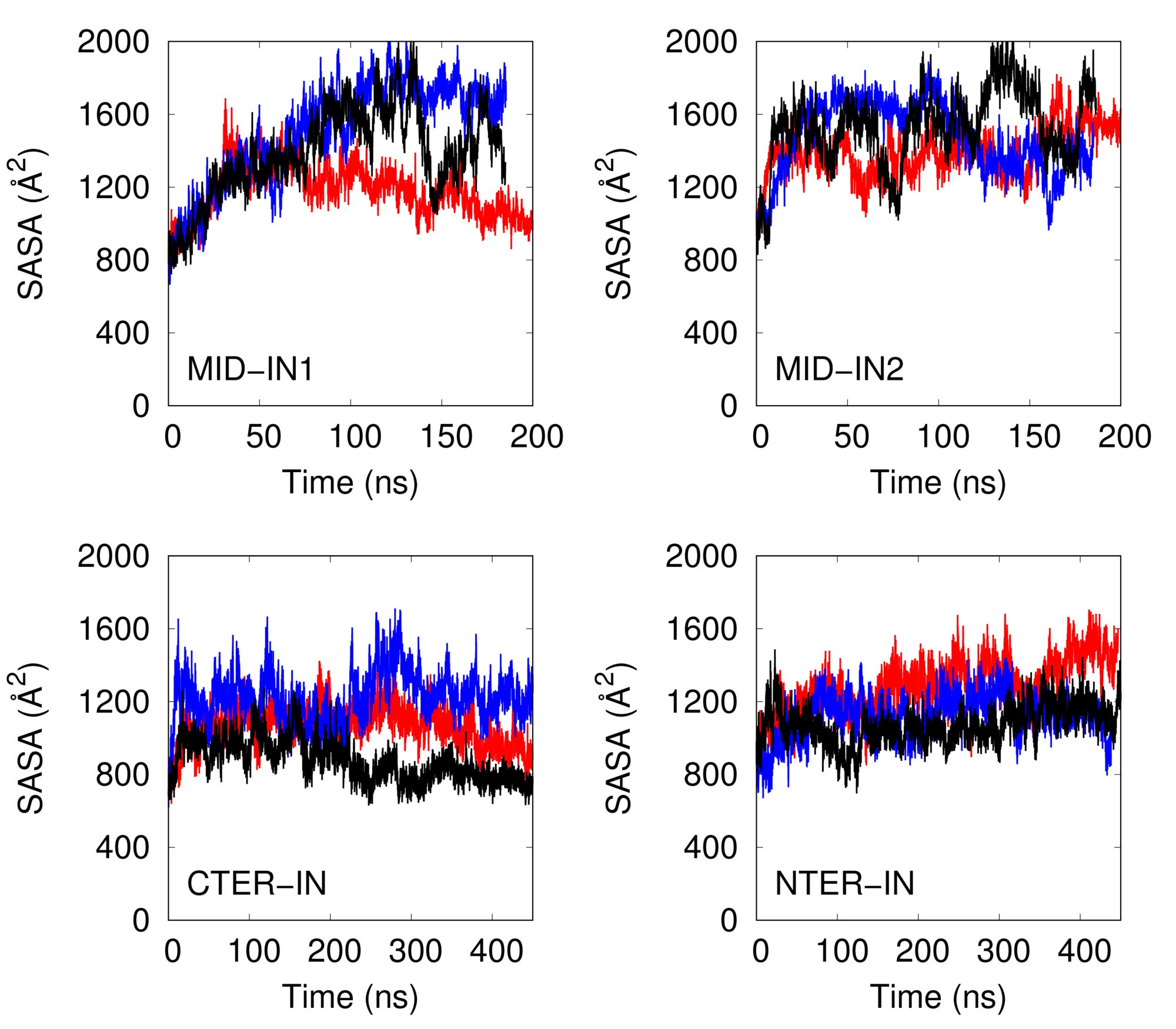

**Figure S2.** SASA of peptides from the MD simulations of MID-IN1, MID-IN2, CTER-IN and NTER-IN modes.

**Table S1.** Occupancy of H-bonds between CXCR4 and *EPI-X4* during the MD simulations

| Donor | Acceptor | Occupancy (%) |
| --- | --- | --- |
| MID-IN1 |  |  |
| *K7-Side* | D187-Side | 33.8 |
| *R3-Side* | D193-Side | 20.7 |
| *L1-Main* | E26-Side | 20.5 |
| *K6-Side* | D262-Side | 19.5 |
| *K6-Side* | E277-Side | 17.6 |
| *S12-Side* | D181-Side | 11.7 |
| MID-IN2 |  |  |
| *K6-Side* | D187-Side | 31.4 |
| *L1-Main* | E32-Side | 28.2 |
| *K7-Side* | D62-Side | 25.5 |
| *Y4-Side* | D97-Side | 21.1 |
| *L1-Main* | E31-Side | 14.6 |
| *Y4-Main* | R30-Main | 12.9 |
| *K7-Side* | E288-Side | 10.0 |
| CTER-IN |  |  |
| *S12-Side* | D262-Side | 35.1 |
| *R3-Side* | D97-Side | 19.8 |
| *T13-Main* | D262-Side | 19.6 |
| *K7-Side* | D187-Side | 14.0 |
| *L1-Main* | D181-Side | 13.8 |
| *T15-Side* | E288-Side | 13.5 |
| *R3-Side* | D187-Side | 12.9 |
| R188-Side | *L16-Side* | 11.4 |
| NTER-IN |  |  |
| *T5-Side* | D187-Side | 69.5 |
| *T5-Main* | D187-Side | 49.9 |
| *R3-Side* | E31-Side | 40.8 |
| *R3-Main* | D97-Side | 37.1 |
| *L1-Main* | D97-Side | 33.8 |
| *K7-Side* | D262-Side | 33.4 |
| R30-Main | *V11-Main* | 26.4 |
| *K6-Side* | R188-Main | 24.3 |
| C28-Main | *T13-Main* | 17.3 |
| *K6-Side* | D187-Side | 17.1 |
| K271-Side | *L16-Side* | 16.2 |
| *R3-Side* | D181-Side | 12.9 |
| R30-Side | *T13-Main* | 12.1 |
| R30-Side | *T13-Side* | 11.7 |
| *T15-Side* | E277-Side | 10.0 |

H-Bond criteria: distance cutoff = 3.0 Å and angle cutoff = 20˚

Residues of EPI-X4 are highlighted in red italics.

**Table S2.** Interaction energies EPI-X4 in different binding modes

| Binding mode | vdW energy (kcal/mol) | Electrostatic energy (kcal/mol) | Total interaction energy (kcal/mol) |
| --- | --- | --- | --- |
| MID-IN1 | -47.3 | -654.8 | -702.1 ± 7.5 |
| MID-IN2 | -49.4 | -682.1 | -731.5 ± 6.3 |
| CTER-IN | -71.4 | -765.7 | -837.1 ± 4.5 |
| NTER-IN | -71.8 | -784.2 | -856.0 ± 3.2 |

The interaction energies were calculated every 0.5 ns from the simulation trajectories under vacuum conditions (i.e., the contributions to the energy by water, membrane and ions were neglected) at the force field level. The errors were estimated by bootstrap analysis using 500 steps.

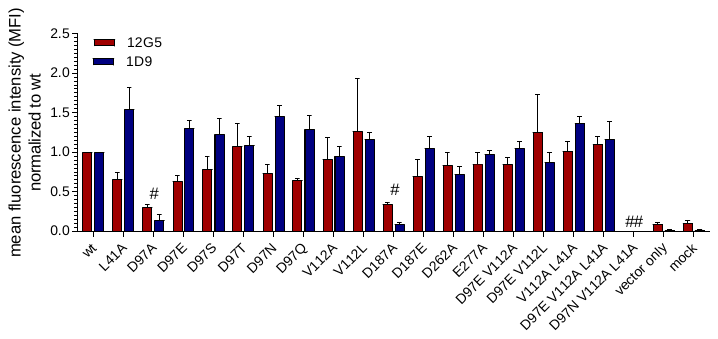

**Figure S3.** CXCR4 expression levels of CXCR4 mutants in 293T cells. Point mutations were introduced into the sequence of CXCR4 in an IRES-GFP expression vector. GPF positive cells were then stained by either the CXCR4 antibody 12G5 (ECL2) or 1D9 (N-terminus) and analyzed by flow cytometry (with the exception of mock control, where also GFP-negative cells were analyzed). Results were normalized to CXCR4 wt expression levels transfected in 293T cells. Shown are data derived from 3 individual experiments ± SEM. # cells were excluded from further binding experiments due to low expression levels, ## no GFP positive cells could be detected.

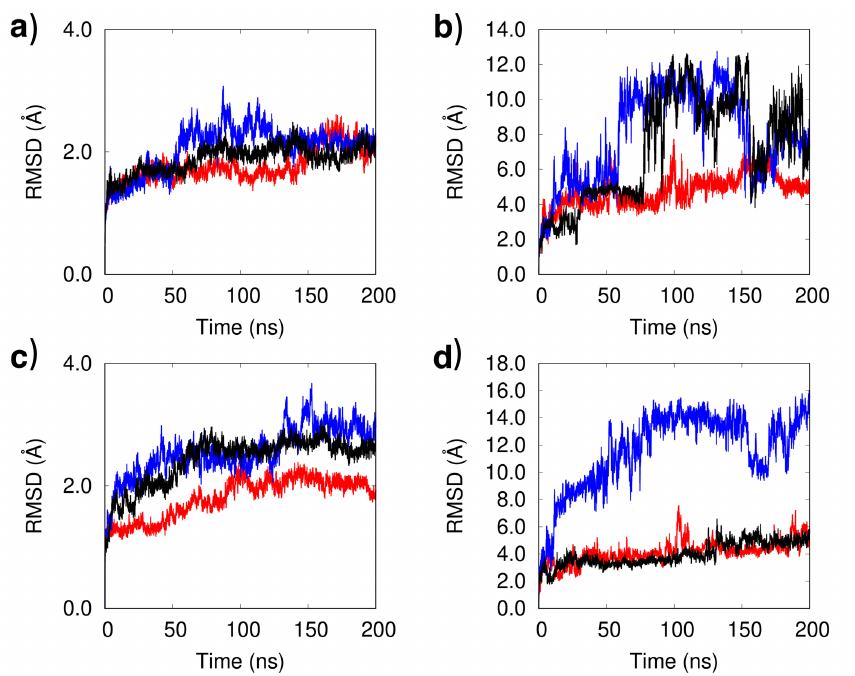

**Figure S4.** RMSD profiles for the CXCR4/WSC02 and CXCR4/JM#21 complexes during the three replicas of the MD simulations. (a) and (b) Backbone RMSD of CXCR4 and WSC02, respectively, in the CXCR4/WSC02 complex. (c) and (d) Backbone RMSD of CXCR4 and JM#21, respectively, in the CXCR4/JM#21 complex.

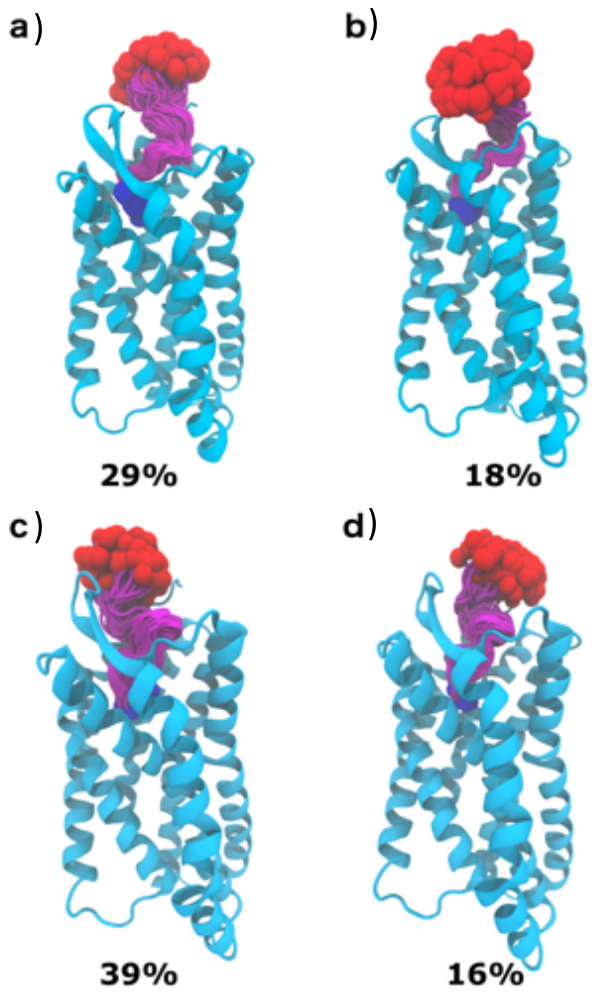

**Figure S5.** Clustering analysis from the MD simulations of CXCR4/WSC02 (a and b) and CXCR4/JM#21 (c and d) complexes. Color scheme is same as that of Figure 1. Two of the top-ranking clusters are shown for each complex. Clustering analysis was performed with the RMSD cutoff of 3 Å, using the VMD plug-in.

**Table S3**. H-bonds formed between protein and peptide in CXCR4/WSC02 and CXCR4/JM#21 complexes

| Donor | Acceptor | Occupancy (%) |
| --- | --- | --- |
| WSC02 |  |  |
| *K7-Side* | E277-Side | 40.8 |
| *K6-Side* | D262-Side | 33.9 |
| *S5-Main* | D187-Side | 32.0 |
| *S5-Side* | D187-Side | 23.9 |
| *R3-Side* | D182-Side | 20.7 |
| *I1-Main* | E288-Side | 17.2 |
| *V2-Main* | D97-Side | 13.2 |
| *R3-Main* | D97-Side | 12.0 |
| *R3-Side* | E2-Side | 11.5 |
| *R3-Side* | D97-Side | 10.2 |
| *I1-Main* | D97-Side | 8.1 |
| JM#21 |  |  |
| *R6-Side* | D262-Side | 81.0 |
| *S5-Side* | D187-Side | 57.6 |
| *S5-Main* | D187-Side | 52.1 |
| *R3-Side* | D97-Side | 38.2 |
| *I1-Main* | E288-Side | 32.4 |
| *K7-Side* | E277-Side | 20.4 |
| *R3-Side* | E31-Side | 18.4 |
| *S12-Side* | D182-Side | 13.0 |
| *R6-Side* | D193-Side | 12.4 |

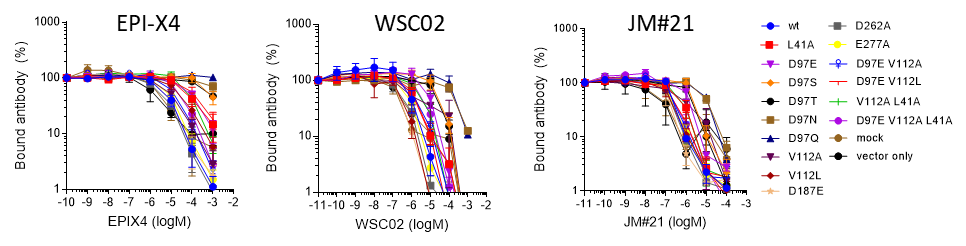

**Figure S6.** EPI-X4, WSC02 and JM#21 interaction to point-mutated CXCR4 by antibody competition. Amino acid substitutions were introduced in the sequence of CXCR4 by site-directed mutagenesis, cloned into an IRES-GFP expression vector and transfected into 293T cells. Afterwards, cells were incubated with serially diluted EPI-X4, WSC02, or JM#21 in presence of a constant concentration of CXCR4 specific antibody (clone 12G5). After 2 hours, bound antibody was analyzed by flow cytometry. Shown are data derived from at least 3 individual experiments ± SEM. (see also Figure 6)

**Table S4.** IC_50_ values for WSC02 and JM#21 determined in an 12G5-competition assay for CXCR4 point mutants

| CXCR4 mutation |  | IC_50_ ± SEM (µM) | |
| --- | --- | --- | --- |
|  |  | WSC02 | JM#21 |
| wt |  | 2.54 ± 1.98 | 0.65 ± 0.10 |
| L41A |  | 1.68 ± 0.78 | 0.72 ± 0.22 |
| D97E |  | 9.62 ± 3.71 | 1.92 ± 0.57 |
| D97S |  | 43.00 ± 23.35 | 5.49 ± 2.26 |
| D97T |  | 34.58 ± 25.75 | 3.91 ± 1.90 |
| D97N |  | 434.53 ± 255.98 | 11.10 ± 2.74 |
| D97Q |  | > 1000 | 23.02 ± 12.89 |
| V112A |  | 0.69 ± 0.24 | 3.48 ± 3.15 |
| V112L |  | 0.48 ± 0.23 | 0.37 ± 0.21 |
| D187E |  | 0.46 ± 0.15 | 0.31 ± 0.24 |
| D262A |  | 0.52 ± 0.26 | 0.23 ± 0.06 |
| E277A |  | 0.77 ± 0.24 | 0.28 ± 0.07 |
| D97E+V112A |  | 1.68 ± 0.34 | 0.46 ± 0.17 |
| D97E+V112L |  | 1.93 ± 0.64 | 0.36 ± 0.16 |
| V112A+L41A |  | 4.73 ± 2.61 | 0.63 ± 0.22 |
| D97E+V112A+L41A |  | 5.80 ± 1.86 | 0.43 ± 0.14 |

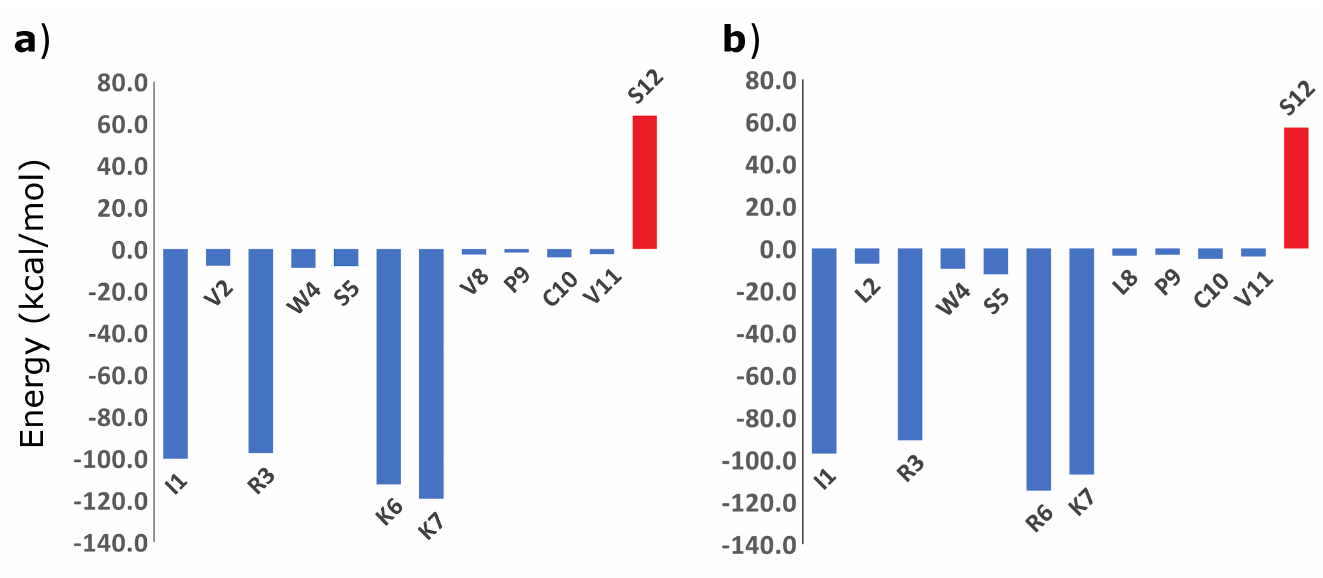

**Figure S7.** Contribution to the interaction energy by individual amino acids of a) WSC02 and b) JM21.

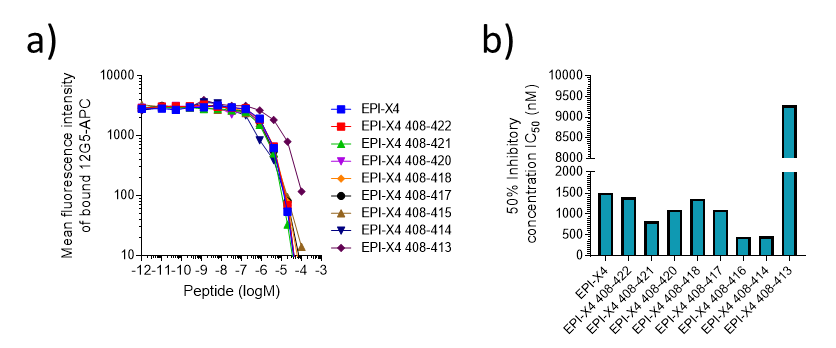

**Figure S8.** C-terminally truncated EPI-X4 competes with an CXCR4 specific antibody. EPI-X4 analogues were designed that are serially truncated at the C-terminus. a) Peptides were serially diluted and added to SupT1 cells together with a constant concentration of a CXCR4 antibody that binds close to the binding pocket of the receptor. After 2 hours, unbound antibody was removed and antibody binding determined by flow cytometry. b) IC_50_ values were determined by non-linear regression. Shown are data derived from one single assay.

**Table S5.** Primers used for site-directed mutagenesis of CXCR4

| **Primer name** | **Primer sequence 5' -> 3'** |
| --- | --- |
| NheI_for | CGGCTAGCATGGAGGGGATCAGTATATACACTTCAG |
| HindIII_elongation_rev | CGAAGCTTTTATTTATCGTATAAAAAAAAGTCTTTTACATCTGTGTTAGCTGGAGTGAAAACTTGAAGACTC |
| 41_Ala_for | TAAAATCTTCGCGCCCACCATCTAC |
| 41_Ala_rev | TTGAAATTAGCATTTTCTTCAC |
| 97_Ser/Glu_rev | AAGGGAAGCGTGATGACAAAGAGG |
| 97_Glu_for | CTGGGCAGTTGAAGCCGTGGCAA |
| 97_Ser_for | CTGGGCAGTTAGTGCCGTGGCAA |
| 97_Thr_for | CTGGGCAGTTACTGCCGTGGCAA |
| 97_Thr_rev | AAGGGAAGCGTGATGACAAAG |
| 97_Asn_for | CTGGGCAGTTAATGCCGTGGCAA |
| 97_Asn/Gln_rev | AAGGGAAGCGTGATGACAAAGAG |
| 97_Gln_for | CTGGGCAGTTCAGGCCGTGGCAA |
| 112_Ala_for | ATGCAAGGCAGCCCATGTCATCT |
| 112_Ala_rev | AGGAAGTTCCCAAAGTACC |
| 112_Leu_for | ATGCAAGGCACTCCATGTCATCTAC |
| 112_Leu_rev | AGGAAGTTCCCAAAGTAC |
| 187_Glu_for | ATATATCTGTGAACGCTTCTACC |
| 187_rev | CTGTCATCTGCCTCACTG |
| 262_Ala_for | GATCAGCATCGCCTCCTTCATCC |
| 262_Ala_rev | CCAATGTAGTAAGGCAGC |
| 277_Ala_for | GTGTGAGTTTGCGAACACTGTGC |
| 277_Ala_rev | CCTTGCTTGATGATTTCC |
| frameshift_for | TAATGAGGGGATCAGTATATACACTTC |
| frameshift_rev | CATGCTAGCCAGCTTGGG |
| stop_for | ACTGAGAAGCTAGACGGACAAG |
| stop_rev | TTCTTCTGGTAACCCATG |

### NMR experiments

#### Methods

To experimentally investigate the conformational properties of EPI-X4 on a medium enriched with negative charges, like the binding pocket of CXCR4, we performed NMR studies of EPI-X4 in the presence of NaOAc. For the NMR experiments with NaOAc, 5 mg of EPI-X4 were dissolved in 450 ml of 10 mM NaP-buffer (NaH_2_PO_4_/Na_2_HPO_4_) and 50 ml D_2_O. For the experiments with acetate ions sodium acetate (C2H3NaO2) was added with a final concentration of 50 mM. If necessary, the pH was adjusted with HCl or NaOH to pH 7. All experiments reported here were recorded on an 850 MHz AVANCE III Bruker system equipped with a 5 mm quadruple resonance QXI 1H/13C/15N/31P probe with a z-gradient. Experiments were carried out at 298 K. Nuclear Overhauser Effect Spectroscopy (NOESY) spectra acquiring 2D homonuclear correlation via dipolar coupling with water suppression using watergate W5 pulse sequence with gradients^1,2^ were recorded for a mixing time of 100, 200 and 300 ms, using 2 x 16k x 256 data matrices, corresponding to acquisition times of ~480 and 8 ms in the t1 and t2 dimensions, respectively. Through-bond connectivity was obtained from a Total Correlation Spectroscopy (TOCSY) spectrum recorded with the MLEV-17 mixing scheme^3^ with water suppression using 3-9-19 pulse sequence with gradients^4,5^, using a 13 µs 90° pulse and a 80 ms mixing period.

NMRFAM-Sparky was used for signal assignment and NOE signal volume determination.^6^ For NOE signal integration a gaussian fit was used with allowing peak motion and adjusting linewidths and baseline fitting. For the 3D structure calculation of EPI-X4 the software package ARIA (Ambiguous Restraints for Iterative Assignment) was used.^7^

#### Analysis of the spectra

Although the linewidths of the NMR spectra were broadened in acidic environment with respect to the neutral medium, no changes in the chemical shift region representing the amide backbone protons were found (Figure S9). This indicates a faster proton exchange event at the peptide backbone compared to EPI-X4 without NaOAc. The remaining protons of the peptide seem to be unaffected. The only affected signal shifted from 3.963 ppm to 3.860 ppm and does not belong to any of the spin systems of EPI-X4 as there is no TOCSY (Total Correlation Spectroscopy) cross-peak at all at this chemical shift. The prominent signal at around 1.8 ppm evidences the presence of acetic acid in the sample. The broader linewidths at the amide backbone region result in the loss of signals at the corresponding region in the TOCSY spectrum (Figure S10). Otherwise, the loss of cross-peaks in the NOESY (Nuclear Overhauser Effect Spectroscopy) spectrum indicates a loss of dipolar spatial couplings between protons of EPI-X4 (Figure S11). Hence, EPI-X4 gains in flexibility and mobility when it is exposed to ions (Figure S12).

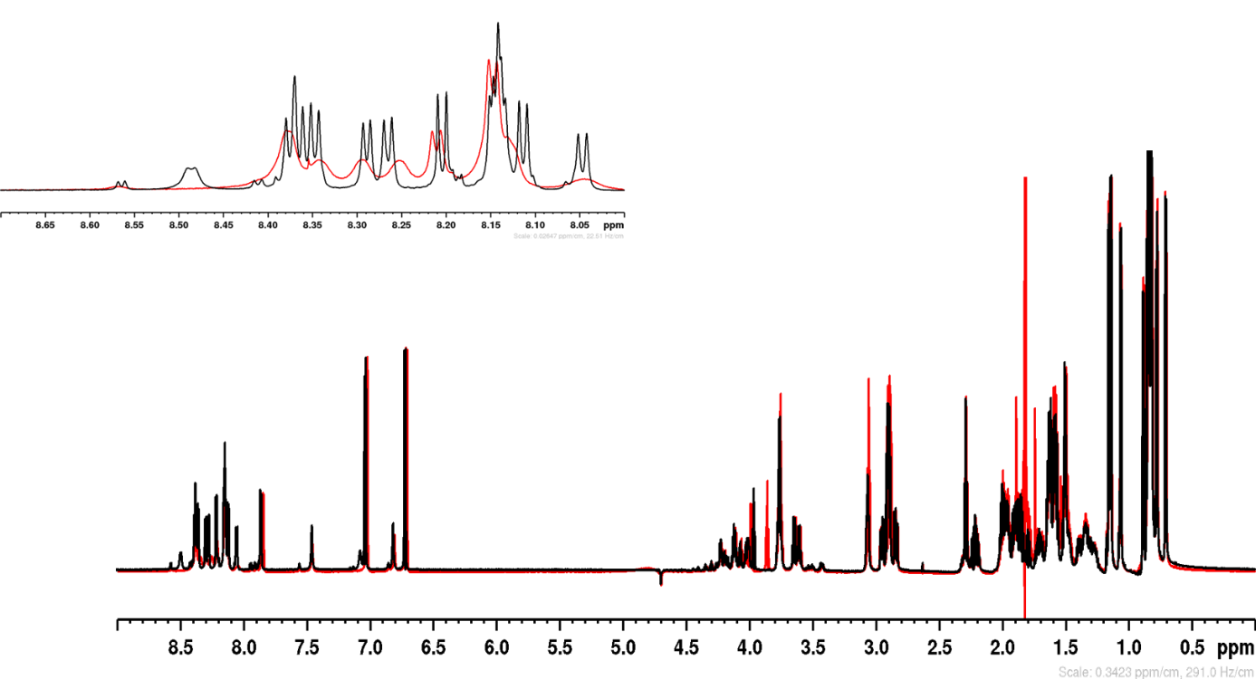

**Figure S9**. ^1^H NMR spectrum of EPI-X4 in NaP-buffer red: with 50 mM NaOAc and black: without NaOAc.

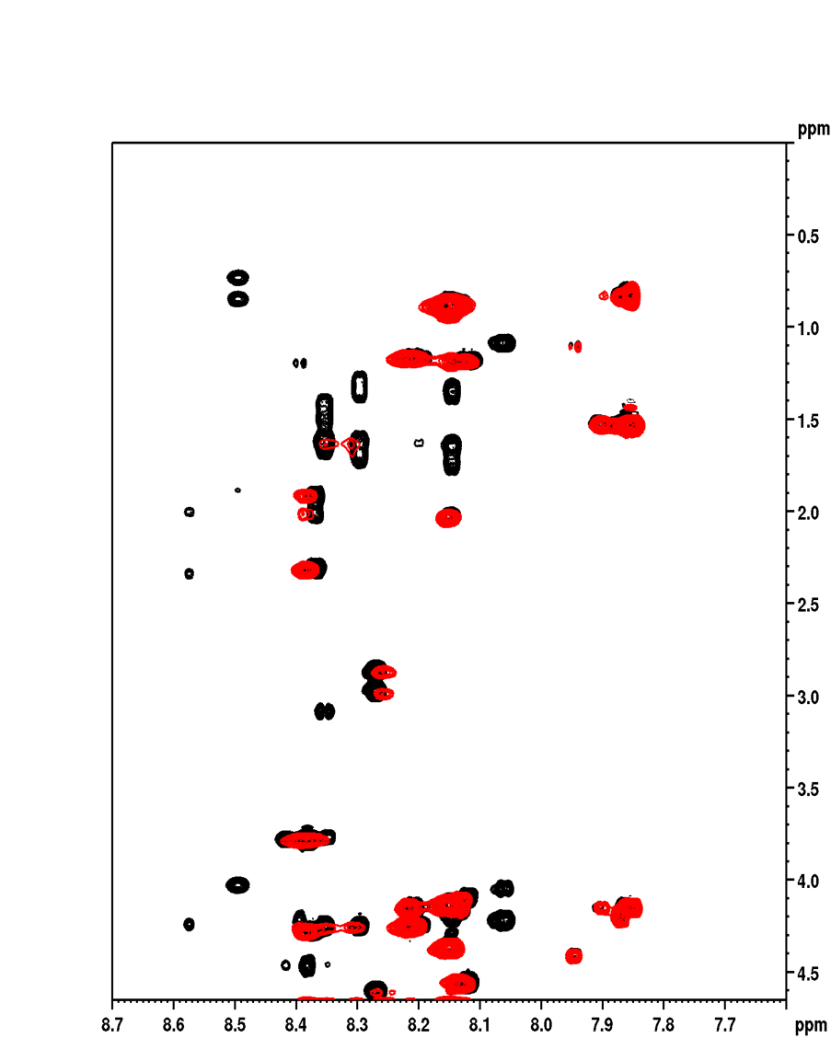

**Figure S10.** ^1^H-^1^H TOCSY NMR spectrum of EPI-X4 in NaP-buffer red: with 50 mM NaOAc and black: without NaOAc

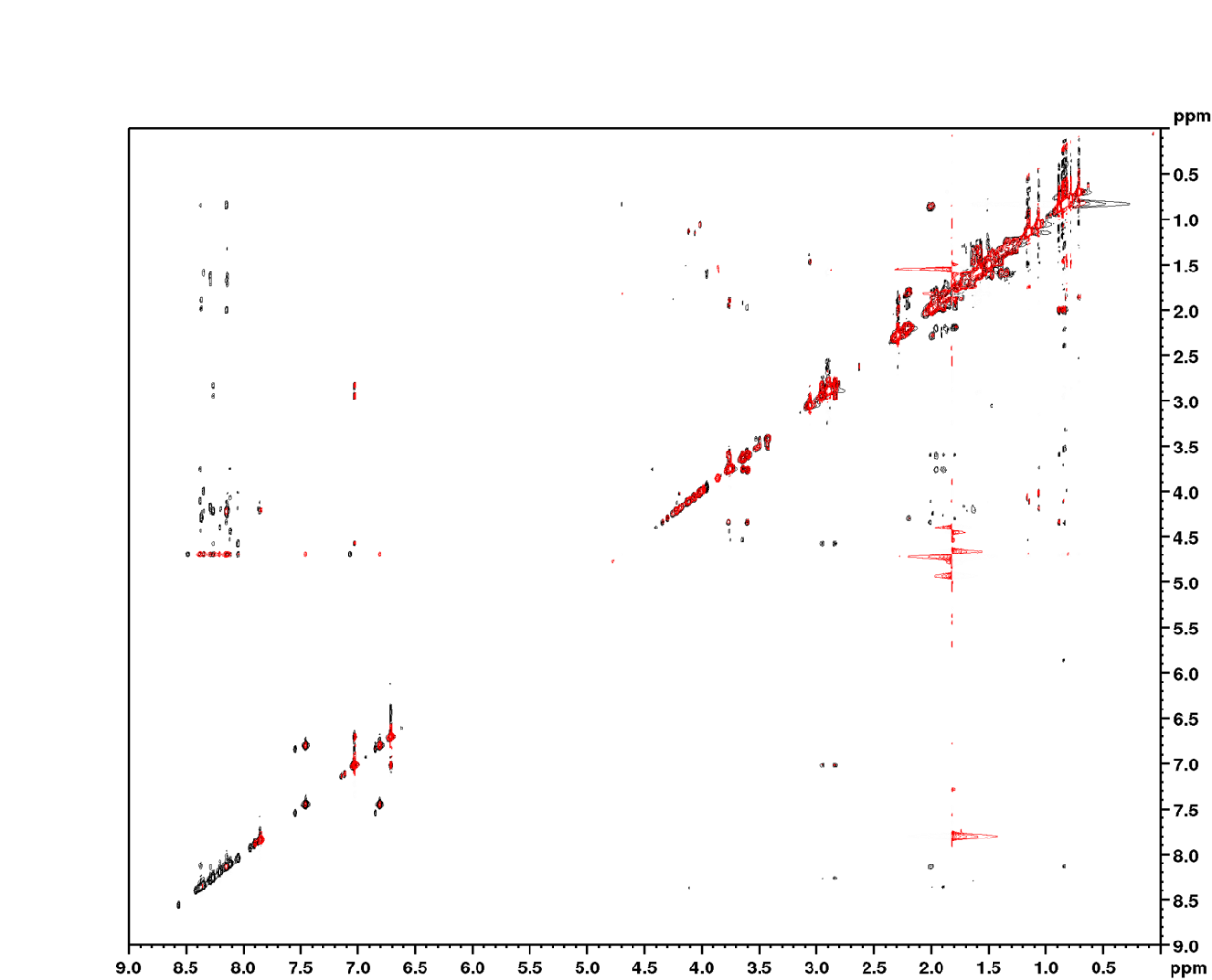

**Figure S11.** ^1^H-^1^H NOESY NMR spectrum of EPI-X4 in NaP-buffer red: with 50 mM NaOAc and black: without NaOAc.

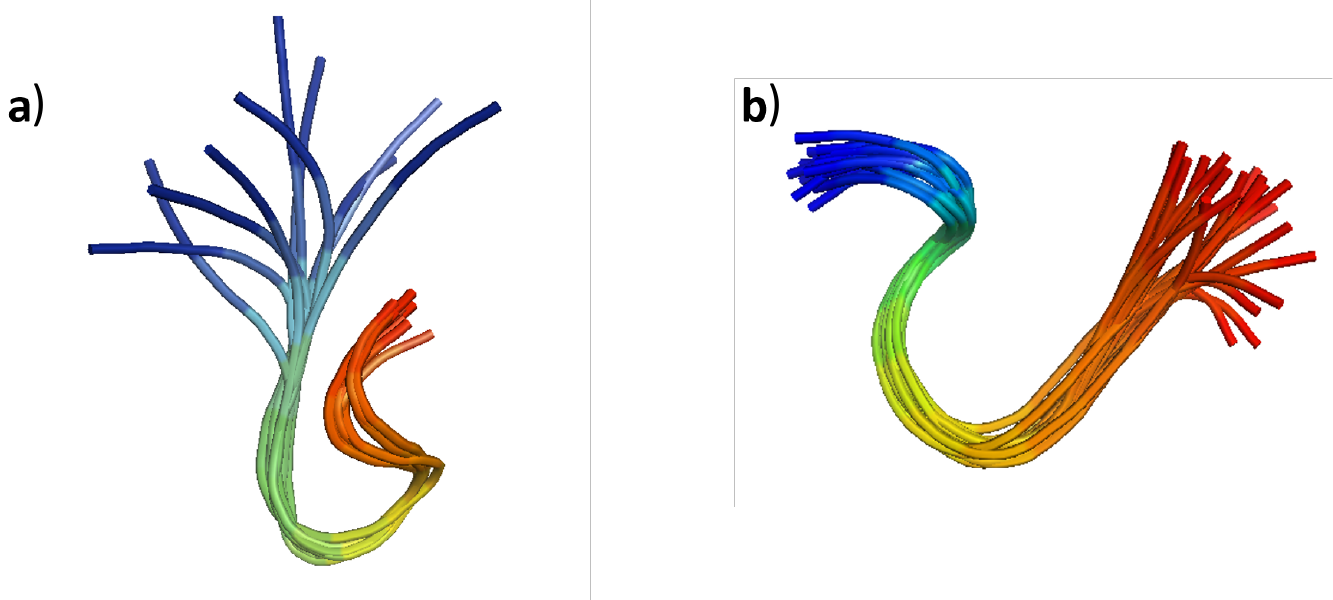

**Figure S12.** EPI-X4 blue to red: N-terminus to C-terminus a) with 50 mM NaOAc b) previously published EPI-X4 without NaOAc. The first 2 amino acids of the N-terminus are free and flexible, while the C-terminus is engaged in hydrogen bonds between Thr6 and Gln10 sidechains as well as a hydrogen bond between the backbone carbonyl of Thr6 and the amide hydrogen of Val11. Additionally, the sidechains of Ser12 and Leu16 are engaged in hydrogen bonds. Arg3 builds a hydrogen bond with its sidechain guanidino group and the carboxyl end of the peptide, when the N-terminus comes close to the C-terminus.

**Table S6.** Criteria used to assess toxicity classes and severity in zebrafish embryos

| **Cytotoxicity / Acute toxicity** | |
| --- | --- |
| L1: | few lysed cells floating in medium; embryos look like wild-type |
| L2: | lysed cells in medium; embryos show some visible tissue damage |
| L3: | embryos show strong tissue damage |
| L4: | embryos are completely disintegrated |
| Nec1: | individual necrotic cells (dark areas in brightfield) |
| Nec2: | many necrotic cells (dark areas in brightfield) |
| **Developmental toxicity** | |
| D1: | developmental delay (slow development) |
| D2: | developmental defects (malformations) |
| **Cardiotoxicity** |  |
| C1: | reduced circulation or heart edema |
| C2: | reduced circulation plus heart edema |
| C3: | no circulation plus heart edema |
| **Neurotoxicity** |  |
| N0: | normal movement in response to touch |
| N1: | reduced movement in response to touch |
| N2: | no movement in response to touch |
| **Overall toxicity (combination of above phenotypes)** | |
| wt: | wild type (no visible phenotype AND normal movement) |
| T1: | embryos that show a phenotype |
| T2: | severe damage so that other phenotypes cannot be assessed (L3, L4, Nec2) |

### References

1.	Jeener, J., et al., *Investigation of Exchange Processes by 2-Dimensional Nmr-Spectroscopy.* Journal of Chemical Physics, 1979. **71**(11): p. 4546-4553.

2.	Liu, M.L., et al., *Improved WATERGATE pulse sequences for solvent suppression in NMR spectroscopy.* Journal of Magnetic Resonance, 1998. **132**(1): p. 125-129.

3.	Bax, A. and D.G. Davis, *MLEV-17-based two-dimensional homonuclear magnetization transfer spectroscopy.* Journal of Magnetic Resonance (1969), 1985. **65**(2): p. 355-360.

4.	Piotto, M., V. Saudek, and V. Sklenar, *Gradient-Tailored Excitation for Single-Quantum Nmr-Spectroscopy of Aqueous-Solutions.* Journal of Biomolecular Nmr, 1992. **2**(6): p. 661-665.

5.	Sklenar, V., et al., *Gradient-Tailored Water Suppression for H-1-N-15 Hsqc Experiments Optimized to Retain Full Sensitivity.* Journal of Magnetic Resonance Series A, 1993. **102**(2): p. 241-245.

6.	Lee, W., M. Tonelli, and J.L. Markley, *NMRFAM-SPARKY: enhanced software for biomolecular NMR spectroscopy.* Bioinformatics, 2014. **31**(8): p. 1325-1327.

7.	Rieping, W., et al., *ARIA2: Automated NOE assignment and data integration in NMR structure calculation.* Bioinformatics, 2007. **23**(3): p. 381-382.
